# The Arabidopsis V-ATPase is localized to the TGN/EE via a seed plant specific motif and acts in a partially redundant manner with the tonoplast enzyme

**DOI:** 10.1101/2020.06.30.179333

**Authors:** Upendo Lupanga, Rachel Röhrich, Jana Askani, Stefan Hilmer, Christiane Kiefer, Melanie Krebs, Takehiko Kanazawa, Takashi Ueda, Karin Schumacher

**Author notes:** Coresponding author Karin Schumacher Heidelberg University Centre for Organismal Studies Dep. Cell Biology Im Neuenheimer Feld 230 69120 Heidelberg Germany. Email of all authors.

## Abstract

Vacuolar-type H^+^-ATPases (V-ATPases) are versatile proton pumps that control the pH of many intracellular compartments in all eukaryotic cells. The localization of the Arabidopsis V-ATPase was previously shown to be determined by the isoforms of subunit a (VHA-a). The incorporation of VHA-a1 targets the V-ATPase to the trans-Golgi network/early endosome (TGN/EE) whilst the incorporation of VHA-a2 or VHA-a3 targets the V-ATPase to the tonoplast. By employing chimeric proteins and site directed mutagenesis we identified a targeting domain (a1-TD) containing an acidic cluster in the N-terminus of VHA-a1 that serves as both an ER export signal and as a TGN retention motif. The a1-TD is conserved among seed plants and we confirmed experimentally that its presence is predictive of TGN/EE- localization. In contrast to many other non-seed plants, the liverwort *Marchantia polymorpha* encodes only a single VHA-a subunit (MpVHA-a) and we show here that it is predominantly localized at the tonoplast. In our attempts to determine if MpVHA-a can functionally replace the Arabidopsis VHA-a isoforms, we used CRISPR/Cas9 to generate null-alleles lacking VHA-a1 and discovered that its function is essential for male gametophyte development but can be replaced by VHA-a2 and VHA-a3 during vegetative growth.

## Introduction

Compartmentalization into distinct membrane-bound organelles that offer different chemical environments optimized for the biological processes that occur within them is a hallmark of eukaryotic cells. Vacuolar-type ATPases (V-ATPases) are rotary engines that couple the energy released by ATP hydrolysis to the transport of protons across membranes of many intracellular compartments. They consist of two subcomplexes: The cytosolic V_1_ subcomplex responsible for ATP hydrolysis composed of eight different subunits (A, B, C, D, E, F, G, H) and the membrane integral V_O_ subcomplex consisting of 6 subunits (a, d, e, c, c’, c”) required for proton translocation (Sze et al., 2002). Although all eukaryotic V-ATPases are strikingly similar regarding their structure and biochemical activity, their biological reach has been greatly diversified by cell-type specific expression and differential subcellular localization (Cotter et al., 2015). Differential targeting of the V-ATPase is mediated by isoforms of subunit a, the largest of the V-ATPase subunits that consists of a C- terminal hydrophobic domain with eight transmembrane domains and a large N-terminal domain that is accessible for cytosolic interaction partners (Zhao et al., 2015; Vasanthakumar et al., 2019). Previous studies in yeast and plants have shown that the targeting information is contained in the N-terminal domain (Dettmer et al., 2006; Kawasaki-Nishi et al., 2001a), however the responsible targeting domain has so far only been addressed for the yeast isoforms Vhp1p and Stv1p. Whereas Vph1p targets the yeast V-ATPase to the vacuole, a motif containing an aromatic residue (WKY) in the N-terminus of Stv1p targets the V-ATPase to the Golgi/endosomal network (Finnigan et al., 2012). Mammals possess four subunit a isoforms (a1, a2, a3, and a4) which steer the V-ATPase to different endomembranes and plasma membranes of specialized cells (Forgac, 2007; Marshansky et al., 2014; Futai et al., 2019). In Arabidopsis AtVHA-a2 (VHA-a2) and AtVHA-a3 (VHA-a3) target the V-ATPase to the tonoplast (Dettmer et al., 2006) where the combined action of V-ATPase and V-PPase energizes secondary active transport and maintains the acidic environment required for the lytic function of plant vacuoles (Sze et al., 1999; Kriegel et al., 2015). AtVHA-a1 (VHA-a1) targets the V-ATPase to the TGN/EE, a highly dynamic organelle that receives and sorts proteins from the endocytic, recycling and secretory pathways (Viotti et al., 2010). The combined action of the V-ATPase and proton-coupled antiporters including the NHX-type cation proton exchangers (Bassil et al., 2011; Dragwidge et al., 2019) and the ClC (Chloride channel) Cl^-^/H^+^ proton antiporters (Fecht-Bartenbach et al., 2007) is responsible for generating and maintaining the acidic pH of the TGN/EE (Luo et al., 2015). Genetic and pharmacological inhibition of the V-ATPase interferes with endocytic and secretory trafficking (Dettmer et al., 2006; Viotti et al., 2010; Luo et al., 2015) and causes defects in both cell division and cell expansion (Dettmer et al., 2006; Brüx et al., 2008). Despite the TGN/EE being the central hub for protein sorting, it is still not clear how the identity of this compartment is specified and how resident proteins required for functionality are maintained while at the same time very similar proteins are sorted and leave the TGN/EE as cargo.

We have shown previously that after assembly involving dedicated ER-chaperones (Neubert et al., 2008), VHA-a1 and VHA-a3 containing Vo-subcomplexes leave the ER via different trafficking routes. Whereas VHA-a1 containing V-ATPases are exported in a COPII (coat protein complex II) - dependent manner (Viotti et al., 2013), VHA-a3 containing complexes have been shown to be delivered in a Golgi-independent manner to the tonoplast (Viotti et al., 2013). Overall, it remains to be addressed how V-ATPases containing the different VHA-a isoforms are sorted in the ER and how they are targeted to their final destinations in the cell. Here, we focus on VHA-a1 to decipher how the V-ATPase is targeted to and retained in the TGN/EE. Targeting of TGN-resident proteins has been shown to be mediated by groups of acidic amino acids (acidic clusters), di-leucine and tyrosine-based motifs in mammals (Bos et al., 1993; Schäfer et al., 1995; Alconada et al., 1996; Xiang et al., 2000). In yeast, retrieval to the late Golgi (equivalent to the TGN) is dependent on the concerted action of aromatic-based amino acid (aa) motifs in the cytoplasmic tails of proteins and slow anterograde transport to the late endosome (Wilcox et al., 1992; Nothwehr et al., 1993; Cooper and Stevens, 1996; Cereghino et al., 1995; Bryant and Stevens, 1997). We have previously shown that the TGN/EE targeting information is contained within the first 228 aa of VHA-a1 (Dettmer et al., 2006). Here, by using chimeric proteins, 3D homology modelling, site-directed mutagenesis and live cell imaging, we identified a region that is required for ER-export as well as TGN/EE-retention. Most plants have multiple VHA-a isoforms, however the VHA-a1 targeting domain (a1-TD) is conserved only among seed plants. The situation in the liverwort *Marchantia polymorpha* with a single gene encoding subunit a (MpVHA-a) that is mostly present at the tonoplast in both Marchantia and Arabidopsis might represent the ancestral state in which acidification of the TGN/EE was achieved by transitory V-ATPase complexes. In support of this notion, we found that MpVHA-a can replace VHA-a2/VHA-a3 at the tonoplast but it fails to rescue the male gametophyte lethality caused by a lack of VHA-a1.

## Results

### Identification of a region in the VHA-a1 N-terminus (a1-TD) that is necessary and sufficient for TGN/EE localization

The tri-peptide motif that mediates Golgi-localization of Stv1p in yeast (Finnigan et al., 2012) is not conserved in VHA-a1 (Supplemental Figure 1) implying that TGN/EE-localization evolved independently. We thus used chimeric proteins consisting of increasing lengths of the cytosolic N-terminal domain of VHA-a1 (a1NT 37, 85, 131, 179 and 228 aa) fused to decreasing lengths of the C-terminal domain of VHA-a3, to further narrow down the region required for TGN/EE targeting. Constructs encoding the chimeric proteins fused to GFP were expressed in Arabidopsis under control of the *UBIQUITIN10* promoter (*UBQ10*; Grefen et al., 2010). Whereas a1NT37a3-GFP, a1NT85a3-GFP and a1NT131a3-GFP all localized to the tonoplast (Supplemental Figure 2), a1NT179a3-GFP and a1NT228a3-GFP were detectable at the tonoplast but also in a punctate pattern reminiscent of the TGN/EE (Figure 1A). TGN/EE localization was confirmed by colocalization with the endocytic tracer FM4-64 (Dettmer et al., 2006) and treatment with the fungal drug Brefeldin A (BFA) that causes the aggregation of the TGN/EE to form BFA compartments (Nebenführ et al., 2002; Dettmer et al., 2006). After 3 hours of BFA treatment, the core of BFA compartments was labelled by a1NT179a3-GFP and a1NT228a3-GFP (Figure 1B). From these observations, we concluded that the region necessary for TGN/EE localization of VHA-a1 (a1-targeting domain; a1-TD) is located between residues L132 and E179.

**Figure 1.**
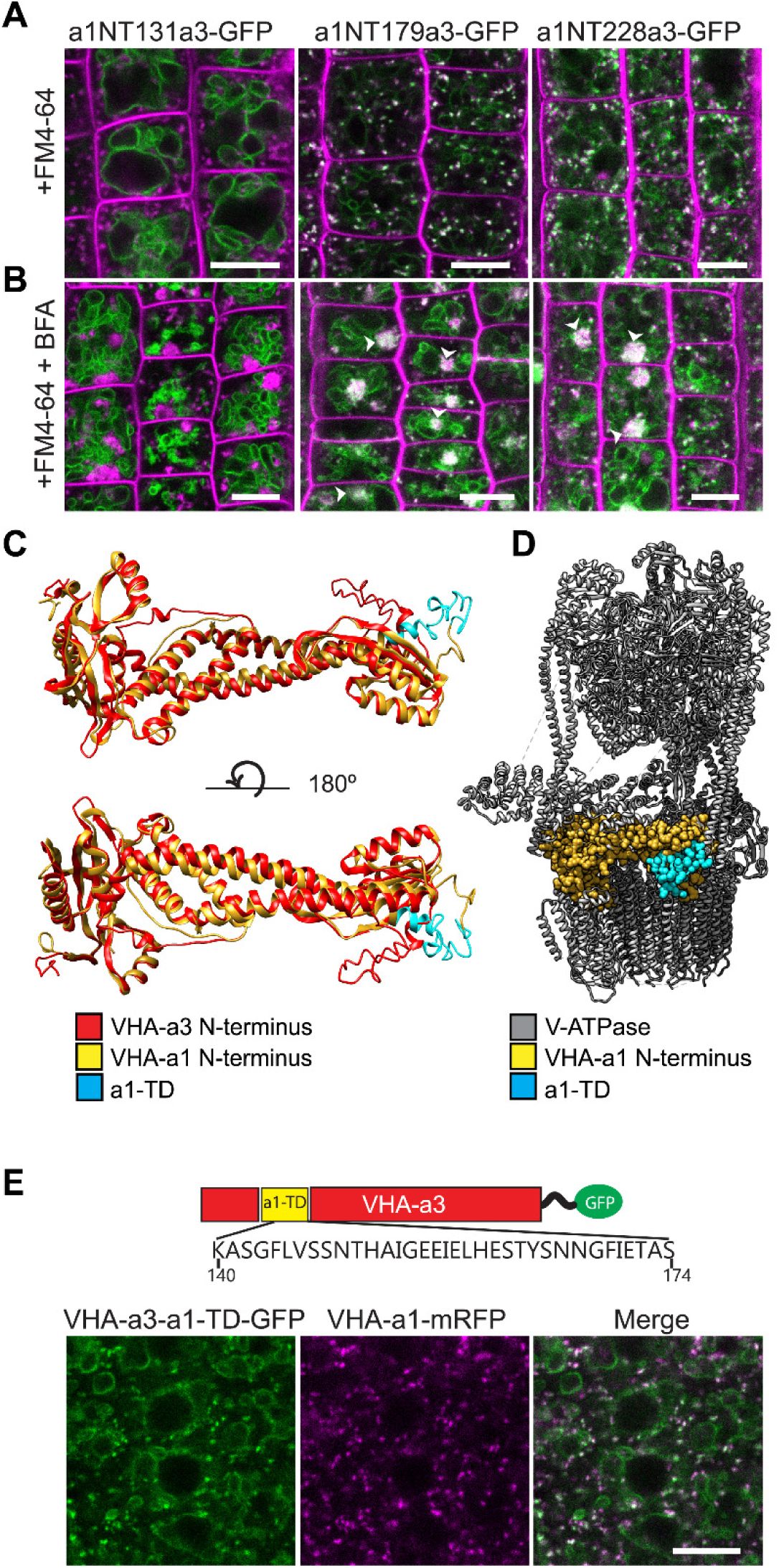
The targeting signal of VHA-a1 is located between L132 and E179 and the a1-TD is sufficient for targeting of VHA-a3 to the TGN/EE. (A) The chimeric constructs a1NT179a3-GFP and a1NT228a3-GFP show dual localization at the TGN/EE and tonoplast. (B) TGN/EE localization of a1NT179a3-GFP and a1NT228a3-GFP was confirmed by treatment of root cells with 50 µM BFA for 3 hours followed by staining with FM4-64 for 20 min. Scale bars = 10 µm. (C) One of the structural differences between the VHA-a1 NT and VHA-a3 NT corresponds to the location of the a1-TD (blue). (D) An alignment of the VHA-a1 NT model with a model of one of the rotational states of the yeast V-ATPase (PDB6O7V) revealed that the VHA-a1 targeting domain is exposed and accessible for recognition. (E) VHA-a3 with the a1-TD (VHA-a3-a1TD-GFP) partially colocalizes with VHA-a1-mRFP. Scale bars = 10 µm.

If the a1-TD contains targeting motifs, it would have to be exposed and accessible when the V-ATPase is fully assembled. To visualize the a1-TD in the three-dimensional (3D) structure, we used I-TASSER (Roy et al., 2010) to generate 3D models of the VHA-a1 and VHA-a3 N-termini with the atomic models of Stv1p and Vph1p from *S. cerevisiae* (Vasanthakumar et al., 2019) serving as the templates, respectively (Figure 1C, Supplemental Table 1 and Supplemental Figure 3). By superimposing the obtained models for VHA-a1 and VHA-a3 N-termini we found that the a1-TD is one of the regions predicted to be different (Figure 1C). To visualize the orientation of the VHA-a1 N-terminus within the V-ATPase complex, the VHA-a1 N-terminus model was aligned to a model of the yeast V-ATPase (PDB6O7V; Vasanthakumar et al., 2019) revealing that the a1-TD would be accessible for recognition in the fully assembled V-ATPase complex (Figure 1D).

To test if the VHA-a1-TD is sufficient to target VHA-a3 to the TGN/EE, we introduced the 34 aa region between K140 and S174 into the N-terminus of VHA-a3 and the resulting GFP fusion construct (UBQ10:VHA-a3-a1-TD-GFP) was transformed into wildtype plants. VHA-a3-a1-TD-GFP was detectable at the tonoplast and at the TGN/EE where it colocalized with VHA-a1-mRFP (Figure 1E). It remains to be determined why TGN/EE localization of VHA-a3-a1-TD is only partial, however it demonstrates that the a1-TD is sufficient to re-route VHA-a3 to the TGN/EE.

### Mutation of the a1-TD in VHA-a1 leads to mislocalization to the tonoplast

Phylogenetic analysis of VHA-a sequences from mono- and dicot species showed that VHA-a1 and VHA-a3 represent two clearly separated clades (Supplemental Figure 4). The region includes a number of amino acids conserved among all of the sequences in the VHA-a1 clade (Figure 2A) but not in the VHA-a3 clade that could either be part of a di-acidic ER-export motif or an acidic cluster involved in TGN/EE-retention. Next, we used site-directed mutagenesis to identify which amino acids are required for correct localization of VHA-a1. Conserved amino acids in the a1-TD were exchanged individually or in combinations of two or three to the corresponding amino acid of VHA-a3. Two mutations had no effect on the localization of VHA-a1 (F134Y and E161S; Figure 2C and D), two mutations led to VHA-a1 being localized to an unknown compartment (E156Q and E156Q + L159T; Figure 2E and F) whereas all other mutations led to a dual TGN/EE and tonoplast localization (L159T, L159T + E161S, E156Q + L159T + E161S (ELE) and E155 + E156 + I157 deletion (ΔEEI); Figure 2G, H, I and J).

**Figure 2:**
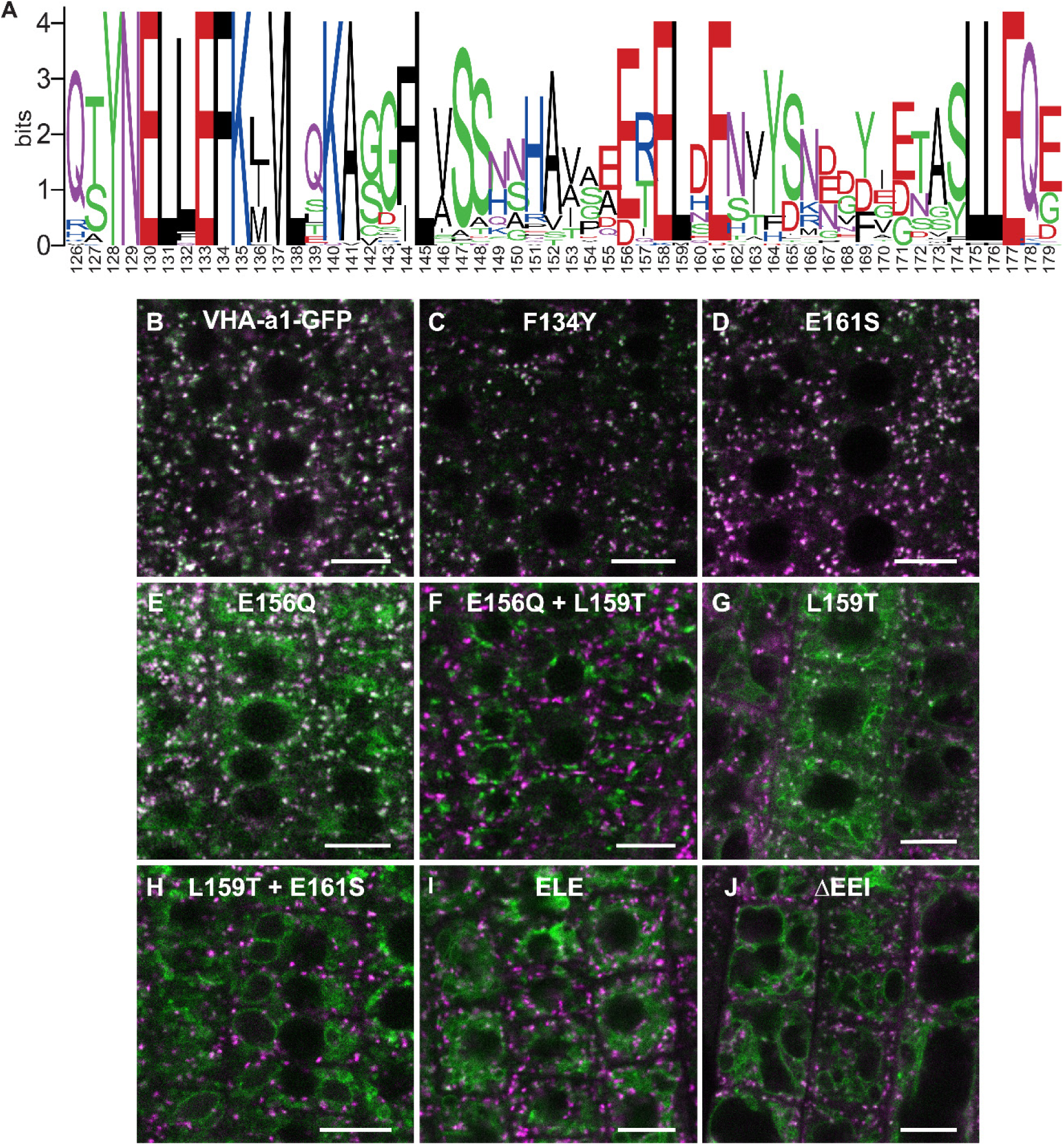
Site directed mutagenesis reveals the importance of conserved amino acids in the targeting of VHA-a1 to the TGN/EE. **(A)** The VHA-a1-clade consensus sequence for the a1-TD region was made on the weblogo platform (Crooks et al., 2004). Sequence numbers are based on Arabidopsis VHA-a1. Conserved amino acids were mutated and their effect on the localization of GFP tagged VHA-a1 was analyzed in the VHA-a1-mRFP background. **(B)** VHA-a1-GFP was co-expressed with VHA-a1-mRFP as a control. The mutations in VHA-a1 produced three classes of punctate patterns that could be classified as TGN/EE only (**C** and **D**), different from VHA-a1-mRFP (**E** and **F**) and TGN/EE and tonoplast (**G**, **H**, **I** and **J**). The overlay of GFP (mutated VHA-a1 proteins) and VHA-a1-mRFP fluorescence is shown. Scale bars = 10 µm.

### The a1-TD is required for COPII-mediated ER-export

As we have shown previously that VHA-a3 containing complexes are delivered in a Golgi-independent manner from the ER to the tonoplast (Viotti et al., 2013), we next asked if the tonoplast signal observed for the mutated VHA-a1 proteins was due to poor retention at the TGN/EE or partial entry into the provacuolar route caused by reduced ER-exit.

To address if the a1-TD contains an ER export motif, we made use of a dominant negative mutation of Sar1 (Sar1-GTP) that has been shown to block COPII-mediated ER-export (daSilva et al., 2004). We expressed AtSar1b-GTP-CFP under a dexamethasone (DEX) inducible promoter (pUBQ10*>GR>AtSar1b-GTP-CFP*; Moore et al., 1998) and used electron microscopy (EM) to confirm that COPII-mediated ER-export was indeed blocked. Upon induction of AtSar1b-GTP-CFP, we observed large clusters of vesicles in the periphery of Golgi stack remnants and ER cisternae appeared swollen (Supplemental Figure 5A). Furthermore, in Arabidopsis transgenic lines expressing the Golgi-marker Sialyl transferase (ST; Boevink et al., 1998) or the brassinosteroid receptor (BRI1; Geldner et al., 2007) as GFP fusion proteins, induction of AtSar1b-GTP-CFP caused their ER-retention and colocalization with AtSar1b-GTP-CFP in bright and often large punctae that might represent the clusters of uncoated COPII-vesicles as observed by EM (Supplemental Figure 5B).

Similarly, the punctate pattern of VHA-a1-GFP disappeared after induction of AtSar1b-GTP-CFP and was replaced by a characteristic ER pattern with dense punctae that colocalized with AtSar1b-GTP-CFP (Figure 3A). In contrast, after the induction of AtSar1b-GTP-CFP, VHA-a3-GFP was not retained in the ER and was exclusively detected at the tonoplast (Figure 3B). To exclude that these differences in ER retention between VHA-a1-GFP and VHA-a3-GFP were due to differences in induction strength of AtSar1b-GTP-CFP, we also blocked ER exit in a line co-expressing VHA-a1-GFP and VHA-a3-mRFP. In cells in which VHA-a1-GFP was retained in the ER, VHA-a3-mRFP was not affected (Figure 3C). Conversely, VHA-a3-a1-TD-GFP is partially retained in the ER upon expression of AtSar1b-GTP-CFP (Figure 3D). Taken together, our data confirms that the ER-export of VHA-a1 is COPII-dependent and reveals that the a1-TD contains an ER-export signal.

**Figure 3:**
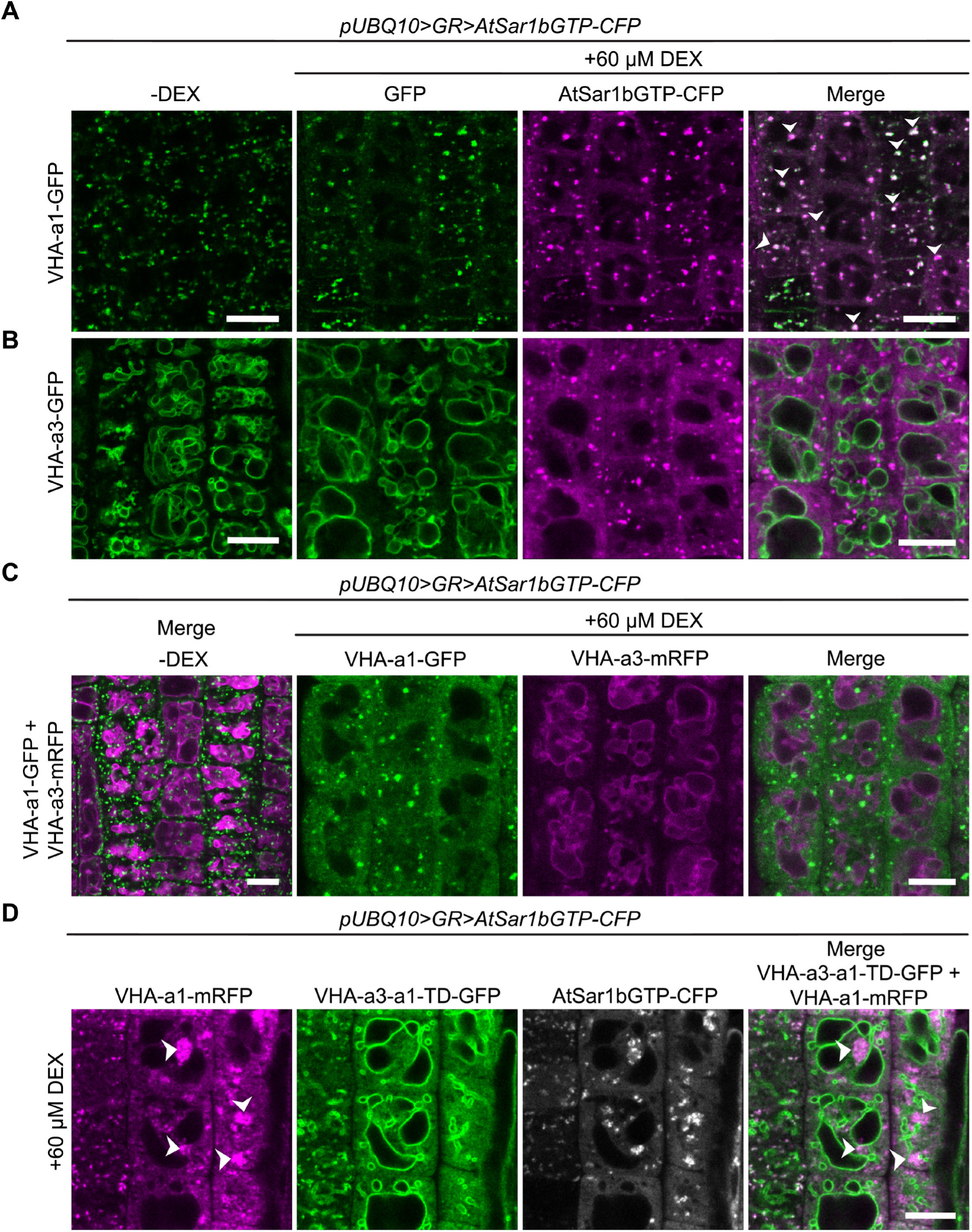
VHA-a1 is retained at the ER after AtSar1b-GTP-CFP expression. After 6 hours of induction with 60 µM DEX, AtSar1b-GTP-CFP is expressed. **(A)** VHA-a1-GFP is retained in the ER and also agglomerates to produce bright punctae which colocalize with AtSar1b-GTP-CFP (white arrows). **(B)** VHA-a3-GFP does not accumulate when exit from the ER via COPII vesicles is blocked by expression of AtSar1b-GTP-CFP. **(C)** When VHA-a1 and VHA-a3 are co-expressed in the same cell only VHA-a1 accumulates in the ER upon induction of AtSar1b-GTP-CFP. **(D)** VHA-a3-a1-TD-GFP is partially retained in the ER after induction of AtSar1b-GTP-CFP. Scale bars = 10 µm.

When AtSar1b-GTP was used to block ER exit, we observed a significant increase in the GFP fluorescence intensity at the tonoplast for all the mutated VHA-a1 proteins (Figure 4A and B) indicating that blocking ER-export causes an increase in the trafficking to the tonoplast via a Golgi-independent route.

**Figure 4:**
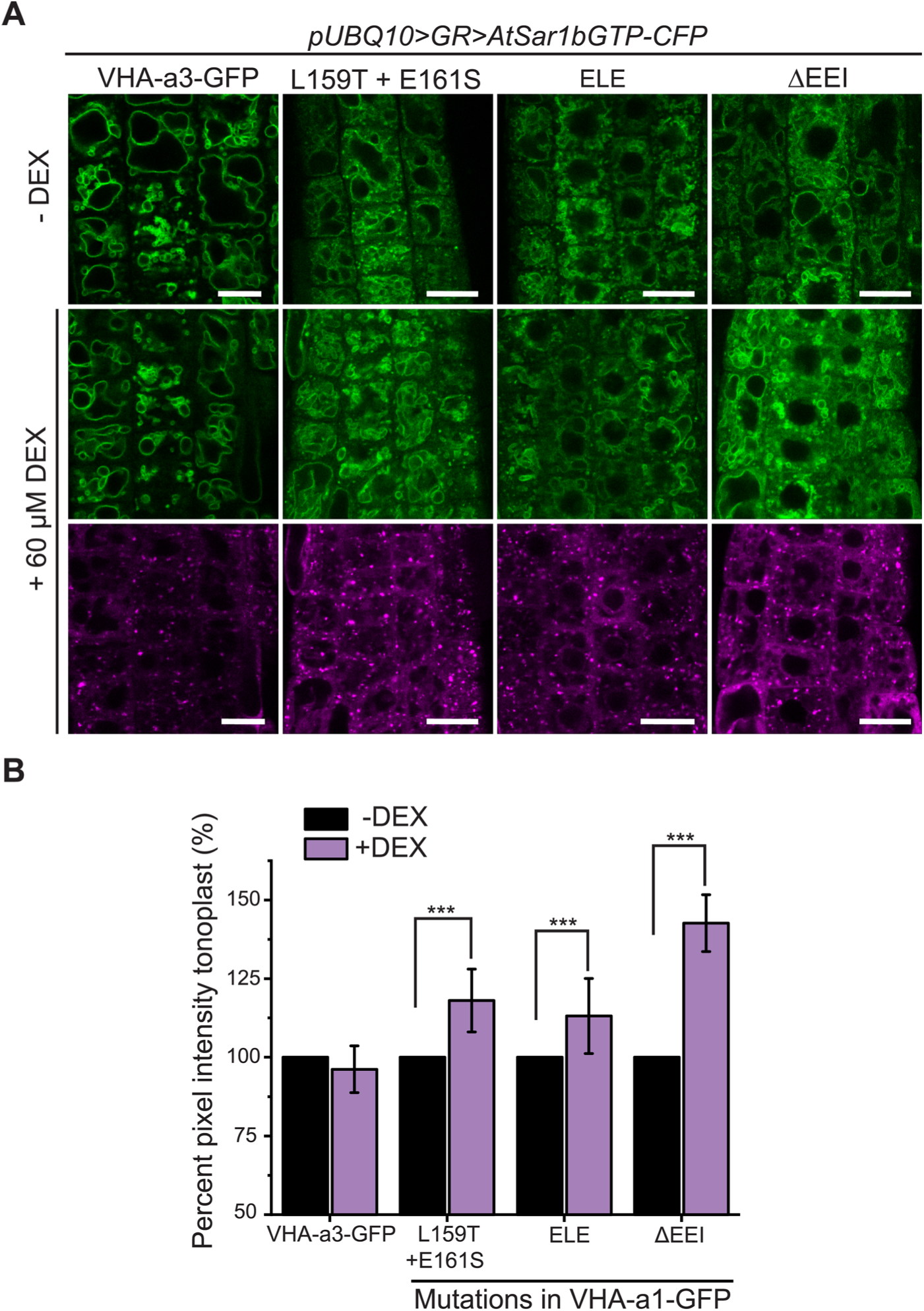
The mutations in the a1-TD affect an ER exit motif. The localization and fluorescence intensity of mutated VHA-a1-GFP proteins was analysed in the presence (+DEX) and absence (-DEX) of AtSar1b-GTP-CFP. **(A)** The mutated VHA-a1 proteins still localized at the tonoplast when exit from the ER via COPII vesicles was blocked by expression of AtSar1b-GTP-CFP. Scale bars = 15 µm. **(B)** The tonoplast fluorescence intensity was measured in the presence and absence of DEX. The fluorescence intensity in the -DEX condition is set to 100% for each genotype. There is a significant increase in the GFP fluorescence intensity at the tonoplast of mutated VHA-a1 proteins when exit from the ER via COPII vesicles is blocked by expression of AtSar1b-GTP-CFP. Error bars indicate SE of n ≥ 216 measurements. Asterisks indicate significant differences between the uninduced and induced conditions (Mann-Whitney test, P < 0.001).

Based on our observation that VHA-a3 carrying a mutation (VHA-a3-R729N-GFP) that renders the complex inactive is retained in the ER when expressed in wildtype but found at the tonoplast in the *vha-a2 vha-a3* mutant (Supplemental Figure 6), we next tested if tonoplast-localization of proteins carrying mutations in the a1-TD increases in the absence of competing complexes containing VHA-a2 and VHA-a3. Indeed, when the mutated VHA-a1 proteins were expressed in the *vha-a2 vha-a3* mutant, the ratio of TGN/EE-to-tonoplast fluorescence intensity increased as compared to the wildtype background (Figure 5A and B). Similarly, VHA-a1 mutants that showed only TGN localization (E161S) or a punctate pattern that is different from VHA-a1-RFP (E156Q and E156Q+ L159T) in the wildtype background displayed a tonoplast signal in the *vha-a2 vha-a3* double mutant (Figure 5A and B). These results strongly support the idea that all mutations affect ER export and that the observed tonoplast signal is not a result of poor retention at the TGN/EE but due to a Golgi-independent pathway to the tonoplast.

**Figure 5:**
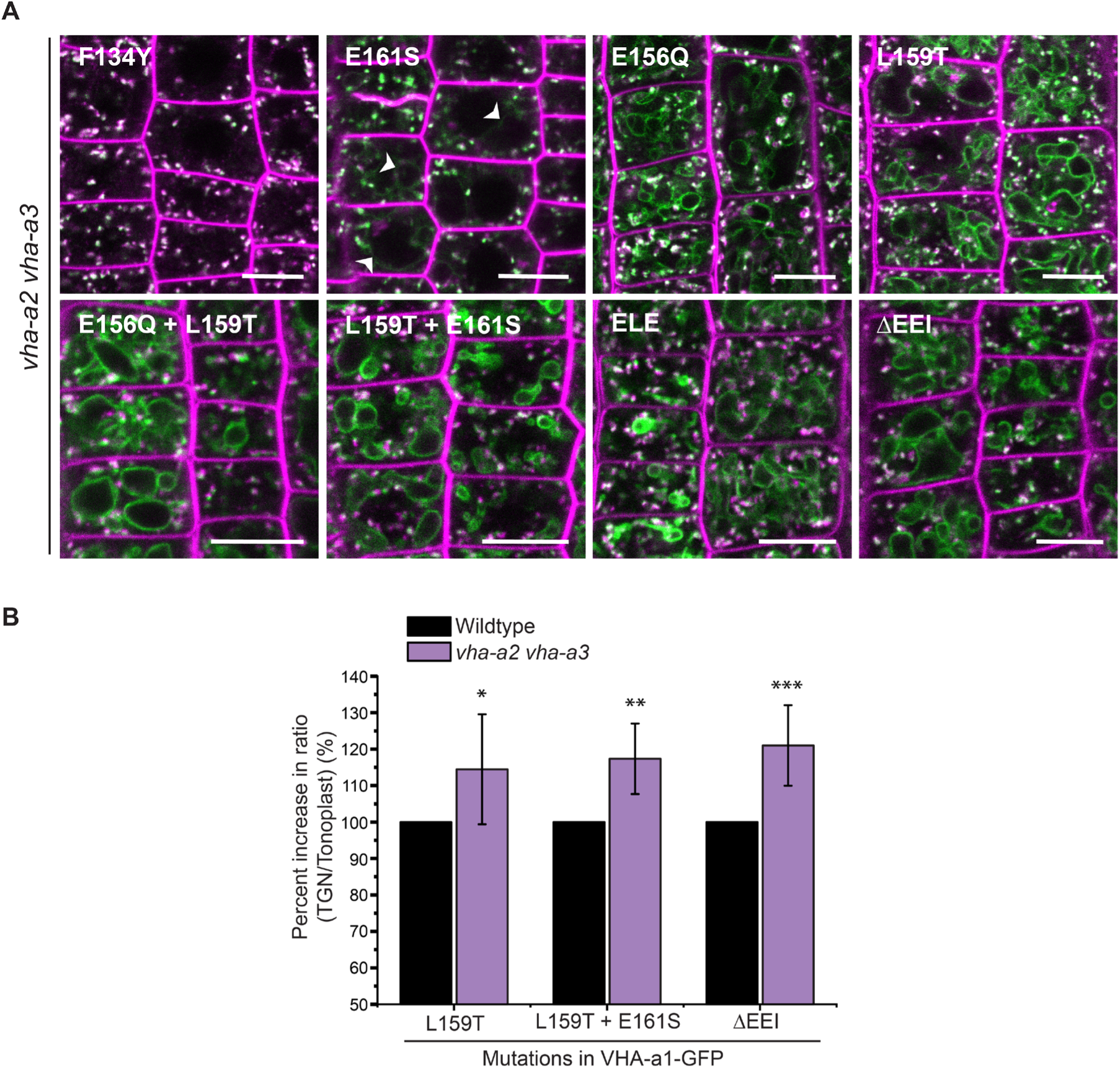
Tonoplast-localization of mutated VHA-a1 increases in the absence of VHA-a2 and VHA-a3. **(A)** The localization of mutated VHA-a1 proteins tagged to GFP were analyzed in the *vha-a2 vha-a3* double mutant background after 20 minutes staining with FM4-64. Scale bars= 10 µm. **(B)** TGN/EE-to-tonoplast fluorescence intensity ratios were calculated for selected mutations in the wildtype background and in the *vha-a2 vha-a3* double mutant background. The ratio of TGN/EE-to-tonoplast fluorescence intensity in the wildtype is set to 100 % for each mutation. The ratio of TGN/EE-to-tonoplast fluorescence intensity of the mutated VHA-a1 proteins is significantly higher in the *vha-a2 vha-a3* double mutant as compared to the wildtype background. Error bars indicate SD of n ≥ 10 ratios calculated from 10 images. Asterisks indicate significant differences between the wildtype and *vha-a2 vha-a3* double mutant ratios (Two-sample *t*-Test, P < 0.05).

### The a1-TD is conserved in the plant kingdom and originates with the Gymnosperms

To trace the evolutionary origin of the a1-TD in the plant kingdom, VHA-a sequences from selected species were subjected to phylogenetic analysis. Our analysis revealed that all VHA-a isoforms from seed plants including the gymnosperm *Pinus taeda* cluster into two distinct clades containing VHA-a1 and VHA-a3 respectively (Supplemental Figure 7 and Figure 7A). Interestingly, the genomes of most basal plants encode multiple isoforms of VHA-a, however they do not fall into either of these two clades implying that duplication of VHA-a occurred independently. Closer analysis of the a1-TD sequence revealed that the hallmarks of the a1-TD (acidic residues flanking a critical leucine residue) are conserved throughout the angiosperms but are absent in bryophytes, lycophytes, ferns and hornworts. Interestingly, the gymnosperm sequence from *P. taeda* contains the a1-TD sequence with the exception of one flanking glutamic residue (Figure 7B). To test if the a1-TD is functionally conserved we generated chimeric proteins consisting of the N-terminal domain of VHA-a from *Pinus taeda* and *Amborella trichopoda* as representatives of the seed plants and *Selaginella moellendorffii* as a non-seed plant fused to the C-terminal domain of VHA-a1 (Figure 7C). Whereas the Pine and Amborella chimeric proteins colocalized with VHA-a1 at the TGN/EE, the Selaginella chimeric protein localized to the tonoplast. These results suggest that the a1-TD domain is conserved within the spermatophytes but not in other green plants.

**Figure 6:**
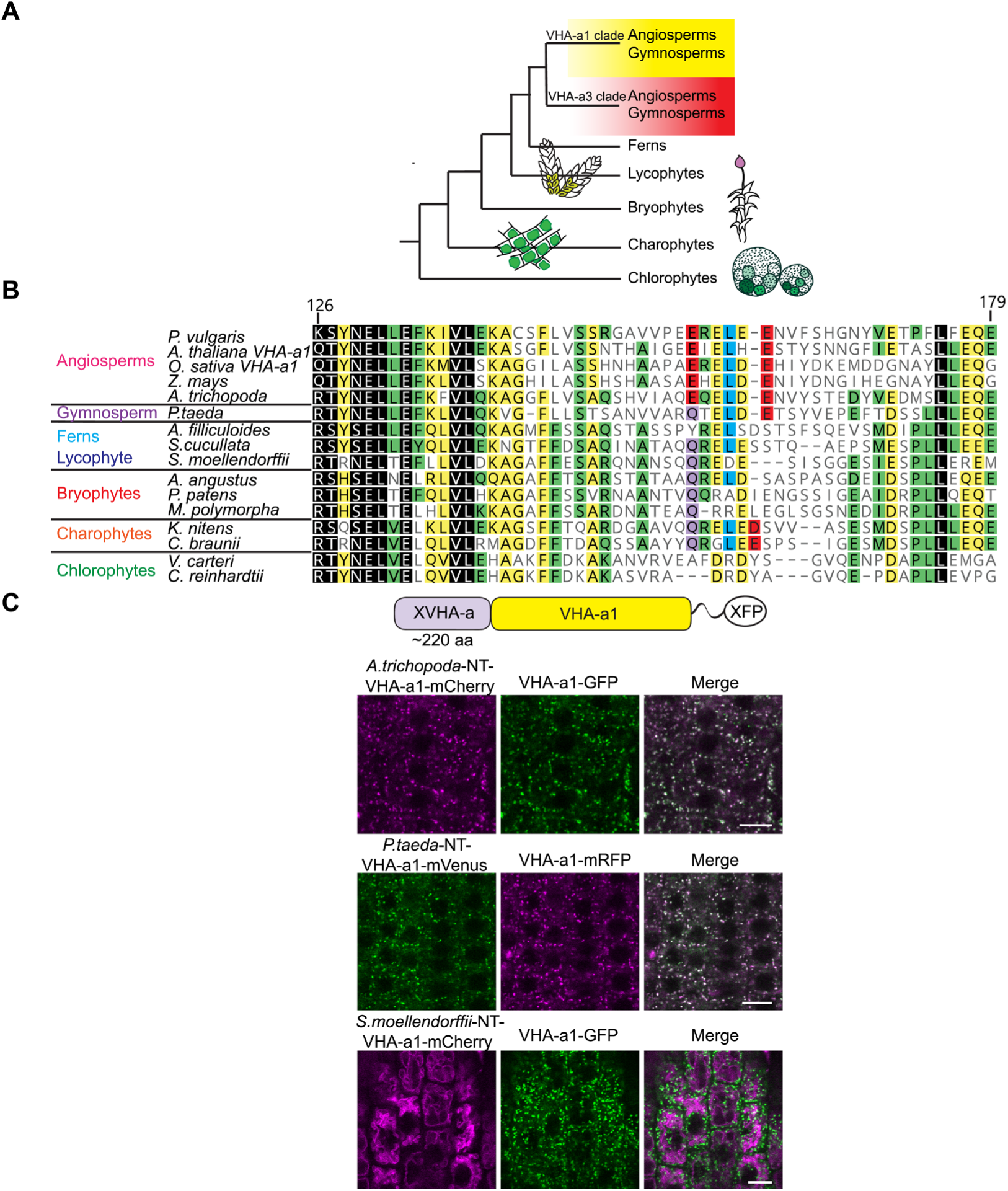
Spermatophyte VHA-a isoforms cluster into two distinct clades, the a1-TD is conserved in the VHA-a1 clade and originates in the gymnosperm sequences. **(A)** Phylogenetic analysis of the N-terminal sequences of VHA-a proteins was done. A graphical summary of the tree is depicted. VHA-a isoforms from seed plants including the gymnosperm *Pinus taeda* cluster into two distinct clades. **(B)** The a1-TD originates in the gymnosperms and it is absent from the chlorophytes, bryophytes, lycophytes and pteridophytes. The sequence of the gymnosperm, *P. taeda* and that of the charophytes contain all the amino acids thought to comprise the ER exit signal with the exception of one flanking glutamic acid residue. The sequence numbers are in reference to the *A. thaliana* VHA-a1 sequence. **(C)** The a1-TD is functionally conserved in gymnosperms and angiosperms. Pine and Amborella chimeric proteins colocalized with VHA-a1 at the TGN/EE. The Selaginella chimeric protein localized to the tonoplast only. Scale bars= 10 µm

**Figure 7:**
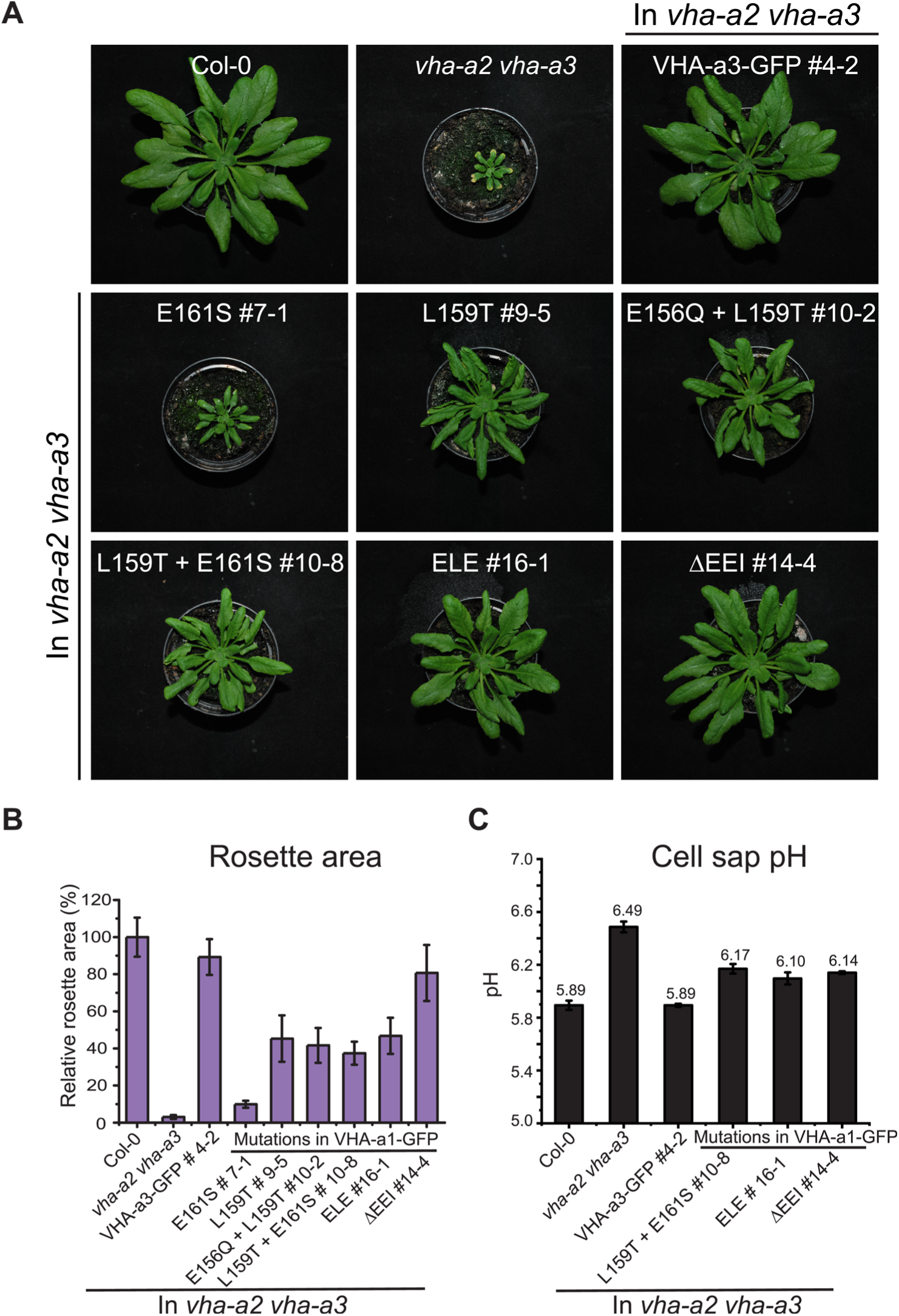
The mutated VHA-a1 subunits complement the *vha-a2 vha-a3* double mutant to varying degrees. Plants were grown in short day conditions (22°C and 10 hours light) for 6 weeks. **(A)** All mutant variants of VHA-a1-GFP displayed bigger rosette size than the *vha-a2 vha-a3* double mutant. E161S which had a faint signal at the tonoplast in the *vha-a2 vha-a3* background also partially complemented the dwarf phenotype of the *vha-a2 vha-a3* double mutant. **(B)** Rosette area of 6- week-old plants grown under short day conditions (n=12). Wildtype rosette size is set to 100 %. VHA-a1 with E155 + E156 + I157 deletion complements the *vha-a2 vha-a3* double mutant the best. **(C)** Mutated VHA-a1 containing V-ATPases have more alkaline cell sap pH values. Error bars represent SD of n = 3 technical replicates.

### Mislocalized VHA-a1 can replace the tonoplast V-ATPase

Mislocalization of VHA-a1 at the tonoplast provides the opportunity to address if the isoforms belonging to the VHA-a1 and VHA-a3 clades are functionally divergent. We used the ability to complement the dwarfed phenotype of the *vha-a2 vha-a3* double mutant (Krebs et al., 2010) in standard long day conditions (Supplemental Figure 8) as well as in short day conditions as (Figure 7A) a proxy. We observed that complementation of the growth phenotype correlated with the intensity of the tonoplast signal with VHA-a1E161S-GFP conferring the lowest and VHA-a1ΔEEI-GFP the highest degree of rescue (Figure 7A and B). The rosette area and diameter of plants expressing VHA-a1ΔEEI-GFP were comparable to that of *vha-a2 vha-a3* double mutant plants expressing VHA-a3-GFP and had similar amounts of protein (Supplemental Figure 9). Three VHA-a1-GFP mutation lines showing high (ΔEEI), intermediate (E156Q + L159T + E161S) and low degrees (L159T + E161S) of rescue were selected for further analysis. Cell sap pH measurements were performed as a proxy of vacuolar pH. Whereas all VHA-a1-GFP mutation lines had more acidic vacuoles than the *vha-a2 vha-a3* mutant none of them reached wildtype vacuolar pH (Figure 7C). Taken together, our data indicates that although VHA-a1 containing complexes are functional when mistargeted to the tonoplast their enzymatic properties are different and they cannot fully replace the vacuolar isoforms.

### The Marchantia V-ATPase is dual localized at the TGN/EE and tonoplast and is functional at the tonoplast in Arabidopsis

The genome of the liverwort *Marchantia polymorpha* encodes a single VHA-a protein allowing us to address if the TGN/EE- or the tonoplast V-ATPase represent the ancestral state or if dual localization of the V-ATPase can also be achieved in the absence of differentially localized isoforms. To examine the subcellular localization of the single, *M*. *polymorpha* VHA-a protein (MpVHA-a), mVenus-tagged MpVHA-a (MpVHA-a-mVenus) was expressed in *M*. *polymorpha* thallus cells. *MpVHA-a-mVenus* driven by the CaMV35S or Mp*EF1α* promoter predominantly localized to the vacuolar membrane, but some additional punctate structures were detectable (Supplemental Figure 10). We then co-expressed MpVHA-a-mVenus and a TGN marker, mRFP-MpSYP6A (Kanazawa et al., 2016), and observed their localization in *M*. *polymorpha* thalli. Again MpVHA-a-mVenus predominantly localized to the vacuolar membrane and partial colocalization with mRFP-MpSYP6A was observed in punctate compartments (Supplemental Figure 10).

Next, we expressed a MpVHA-a-mVenus fusion construct (*UBQ10:MpVHA-a-mVenus*) in the Arabidopsis wildtype background. Whereas MpVHA-a-mVenus localization at the tonoplast of Arabidopsis root cells was clearly visible (Supplemental Figure 11A), a clear punctate pattern was not observed. However, after BFA treatment, the core of BFA compartments was labelled with MpVHA-a-mVenus (Supplemental Figure 11B) indicating that MpVHA-a is also present at the TGN/EE. To test for functionality, MpVHA-a-mVenus was expressed in the *vha-a2 vha-a3* mutant background. MpVHA-a-mVenus localized to the TGN/EE and tonoplast and was found in the core of FM4-64 labelled BFA compartments (Figure 8A). A growth assay conducted in short day conditions (SD; 22°C and 10 hours light) revealed that V-ATPases that incorporate MpVHA-a can complement the *vha-a2 vha-a3* double mutant to wildtype levels (Figure 8B).

**Figure 8.**
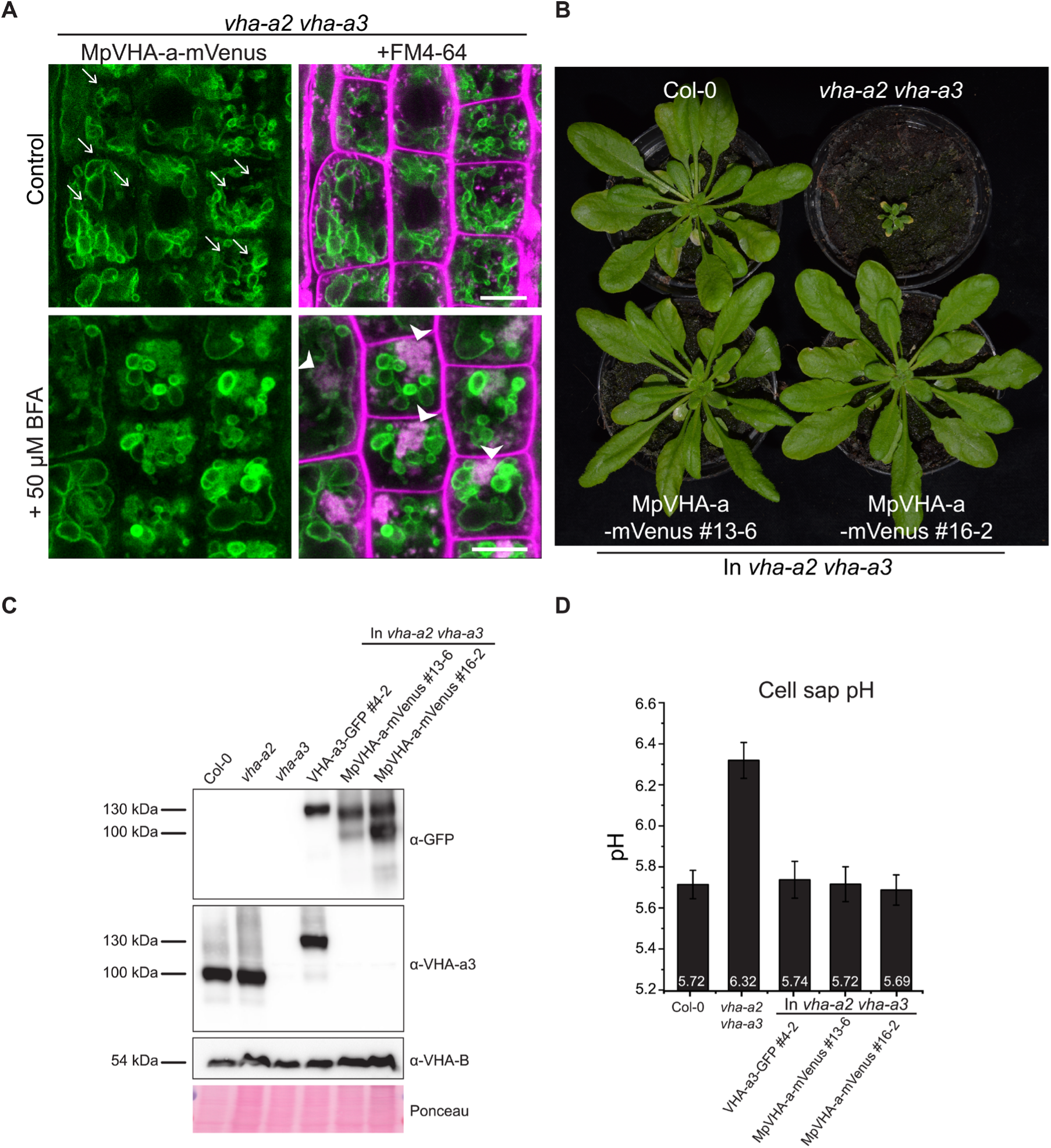
MpVHA-a containing complexes are functional in Arabidopsis and can replace VHA-a3 and VHA-a2 at the tonoplast. **(A)** MpVHA-a-mvenus is dual localized at the tonoplast and TGN/EE in the *vha-a2 vha-a3* background. The TGN/EE localization was confirmed by treatment of root cells with 50 µm BFA for 3 hours followed by staining with FM4-64 for 20 min. The core of BFA compartments were labelled with MpVHA-a-mvenus and FM4-64. Scale bars = 10 µm. **(B)** MpVHA-a-mVenus can also fully complement the dwarf phenotype of the *vha-a2 vha-a3* double mutant. Plants for complementation assay were grown in SD conditions for 6 weeks. **(C)** Western blot of tonoplast membrane proteins from 6-week-old rosettes. Protein levels in the MpVHA-a-mVenus complementation lines is comparable to the VHA-a3-GFP complementation line. **(D)** Cell sap pH of rosette leaves from plants grown in LD conditions for 3 weeks. MpVHA-a-mVenus complexes restore the cell sap pH of the *vha-a2 vha-a3* double mutant to wildtype levels. Error bars represent SD of n = 2 biological replicates.

### VHA-a1 has a unique and essential function during pollen development which cannot be fulfilled by VHA-a2, VHA-a3 or MpVHA-a

Next, we wanted to test if MpVHA-a can also replace VHA-a1 at the TGN/EE. Null alleles of single copy encoded VHA-subunits cause male gametophyte lethality (Dettmer et al., 2005) and it has thus not been possible to identify homozygous T-DNA mutants for VHA-a1. To avoid T-DNA related silencing problems in complementation experiments with heterozygous *vha-a1/+* mutants, we used CRISPR/Cas9 under control of an egg cell specific promoter to generate mutant alleles (Wang et al., 2015). Different regions of *VHA-a1* were targeted using four different guide RNAs (gRNAs 1-4, Supplemental Figure 12A). Wildtype plants as well as *VHA-a1-GFP* expressing transgenic plants were transformed and T1 plants were analyzed by sequencing of PCR products spanning the CRISPR site. Four alleles were selected for further studies: *vha-a1-1* containing a 260 bp deletion which eliminates the start codon and *vha-a1-2*,*-3* and *-4* that each contain single base pair insertions leading to frameshifts and early stop codons (Supplemental Figure 12A).

Surprisingly, we did not only identify heterozygous *vha-a1/+* but also homozygous and bi-allelic *vha-a1* individuals (Supplemental Table 2). The latter two were indistinguishable from wildtype during vegetative growth, but failed to produce seeds due to a defect in pollen development (Figure 9A). To confirm that the defect in pollen development is indeed caused by a lack of VHA-a1, we analysed if it is rescued by *VHA-a1-GFP*. Although *VHA-a1:VHA-a1-GFP* (Dettmer et al., 2006) is also targeted by the chosen gRNAs and *UBQ10:VHA-a1-GFP* is targeted by gRNAs 2,3 and 4, the use of appropriate primer combinations allowed us to distinguish between mutations in the endogenous locus and the transgene (Supplemental Figure 12B). Plants that carried mutations corresponding to *vha-a1-2* in the transgene did not express VHA-a1-GFP confirming that it is indeed a null allele (Supplemental Figure 12C). Importantly, all plants with no wildtype allele of either *VHA-a1* or *VHA-a1-GFP* were defective in pollen development, whereas *vha-a1* mutants expressing VHA-a1-GFP under the control of *UBQ10* promoter showed normal pollen development (Figure 9A).

**Figure 9.**
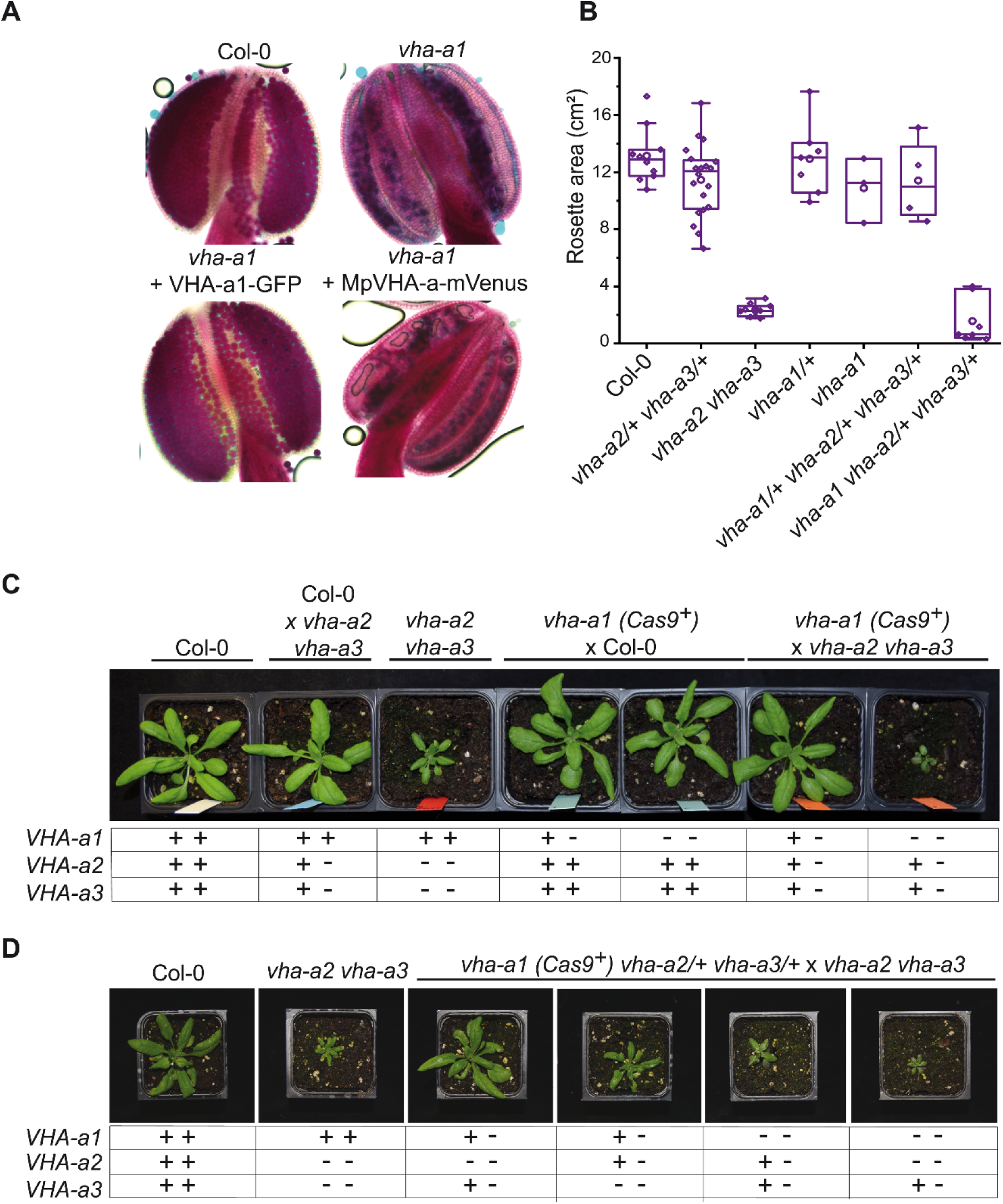
VHA-a1 is essential for pollen development in which it cannot be replaced by VHA-a2, VHA-a3 and MpVHA-a. **(A)** Misshaped microspores/pollen in *vha-a1* anthers were visualized using Alexander’s stain. VHA-a1-GFP rescues the pollen phenotype of *vha-a1*, while MpVHA-a-mVenus does not. **(B, C)** *vha-a1 (Cas9^+^)* was crossed with the *vha-a2 vha-a3* double mutant. Analysis of F1 plants revealed that *vha-a1 vha-a2/+ vha-a3/+* is reduced in growth. Rosette areas of 3.5-week-old plants were quantified. **(D)** *vha-a1 (Cas9^+^) vha-a2/+ vha-a3/+* was crossed with the *vha-a2 vha-a3* double mutant. *vha-a1/+ vha-a2/+ vha-a3* is smaller than *vha-a1/+ vha-a2 vha-a3/+* and *vha-a1 vha-a2 vha-a3/+* was the smallest mutant found. 4-week-old plants are shown. Plants were grown under long day conditions (22°C and 16 hours light).

We next established stable lines expressing *VHA-a1-GFP in the vha-a1-1* background and performed reciprocal crosses to determine if transmission via the female gametophyte is also affected. As expected, *vha-a1-1* was only transmitted via the male gametophyte in the presence of *VHA-a1-GFP*. In contrast, transmission of *vha-a1-1* via the female gametophyte did not require the presence of the transgene indicating that development of the female gametophyte is not affected (Supplemental Table 3).

As homozygous *vha-a1* mutants cannot be transformed, rescue experiments would require crosses with a pollen donor that carries a wildtype allele of *VHA-a1* so that complementation could only be determined in the F2. For crosses we thus used *vha-a1* mutants that still contained the CRISPR T-DNA *(Cas9^+^)* so that the incoming wildtype allele would be mutated and homozygous *vha-a1* mutants could be obtained in the resulting F1. Using this strategy, we identified bi-allelic and homozygous *vha-a1* mutants expressing *UBQ10:MpVHA-a-mVenus* and found that it was localized both at the TGN/EE and the tonoplast (Supplemental Figure 13). However, MpVHA-a-mVenus did not rescue the pollen phenotype of *vha-a1* (Figure 9A) indicating that the TGN/EE-localized V-ATPase of seed plants has acquired a unique function during pollen development.

### During vegetative growth V-ATPases containing VHA-a2 and VHA-a3 compensate for the lack of VHA-a1

Given the importance of acidification of the TGN/EE for endomembrane trafficking and the fact that we have previously reported that RNA-mediated knock-down of VHA-a1 causes reduced cell expansion (Brüx et al., 2008), the observation that the *vha-a1* mutant displays no different phenotypes from wildtype during vegetative growth is unexpected. However, we observed that *vha-a1* root cells are hypersensitive to the V-ATPase inhibitor Concanamycin A (ConcA) in contrast to *vha-a1* roots expressing *UBQ10:VHA-a1-GFP*. This result indicates that a target of ConcA is present at the TGN/EE in *vha-a1* roots albeit in small amounts or with a higher sensitivity to ConcA (Supplemental Figure 14A). To test if VHA-a2 or VHA-a3-containing V-ATPases might compensate for the lack of VHA-a1, *vha-a1-1 (Cas9^+^)* was crossed with the *vha-a2 vha-a3* double mutant. *vha-a1 vha-a2/+ vha-a3/+* mutants obtained from this cross were significantly smaller than *vha-a2/+ vha-a3/+* individuals indicating that tonoplast V-ATPases might indeed compensate for the lack of VHA-a1 during vegetative growth (Figure 9B and C). Subsequently, *vha-a1 (Cas9^+^)* was crossed with *vha-a2* and *vha-a3* single mutants to determine if both are able to compensate for the lack of VHA-a1. *vha-a1 vha-a2/+ and vha-a1 vha-a3/+* plants both had reduced rosette sizes, however the latter showed a stronger reduction which is in accordance with VHA-a3 being expressed at higher levels than VHA-a2 (Supplemental Figure 14B). To analyze mutants with even lower numbers of *VHA-a* wildtype alleles *vha-a1 (Cas9^+^) vha-a2/+ vha-a3/+* was crossed with the *vha-a2 vha-a3* double mutant. Consistent with the results of the single mutant crosses, *vha-a1/+ vha-a2/+ vha-a3* was smaller than *vha-a1/+ vha-a2 vha-a3/+.* The smallest plants obtained were *vha-a1 vha-a2 vha-a3/+* (Figure 9D). Neither we found *vha-a1 vha-a2/+ vha-a3* nor the homozygous triple mutant *vha-a1 vha-a2 vha-a3*, suggesting that they are not viable.

## Discussion

### The information needed to deliver and keep the V-ATPase at the TGN/EE is contained in the a1-TD

The compartments of the eukaryotic endomembrane system are acidified to varying degrees by the activity of the vacuolar H^+^-ATPase (V-ATPase). Acidification enables specific biochemical reactions as well as secondary active transport and can thus be considered a central component of compartmentation. Throughout eukaryotes, isoforms of the membrane-integral V_O_-subunit a are used to achieve differential targeting of the V-ATPase and although we know very little about the underlying mechanisms, it is clear that differential targeting has arisen independently in different groups. Whereas the targeting information is contained in the N-terminus of the Golgi-localized yeast isoform Stv1p (Finnigan et al., 2012), it was shown to be provided by the C-terminal part for some of the 17 isoforms of *Paramecium tetraurelia* that localize the V-ATPase to at least seven different compartments (Wassmer et al., 2006). Moreover, the targeting motif identified in Stv1p (Finnigan et al., 2012) is neither conserved in plant nor mammalian subunit a isoforms implying that differential targeting mediated by subunit a evolved independently in the different eukaryotic lineages. Given the essential role that the V-ATPase plays in endocytic and secretory trafficking (Dettmer et al., 2006; Luo et al., 2015), we set out to identify the mechanism underlying its highly specific localization at the TGN/EE (Dettmer et al., 2006). Based on our previous studies, in which we had shown that the targeting information is located within the 228 amino acids of VHA-a1 (Dettmer et al., 2006), we first narrowed down this region to a 30 aa region (a1-TD) which we show here to be both necessary and sufficient for TGN/EE-localization. Structural modeling revealed that the a1-TD that contains several highly conserved acidic amino acids is exposed and accessible for interaction partners. Acidic clusters have been shown to be involved in TGN-localization (Schäfer et al., 1995; Alconada et al., 1996; Xiang et al., 2000) and we thus used site-directed mutagenesis and found that mutation of conserved aa in the a1-TD indeed leads to partial mislocalization of VHA-a1 to the tonoplast as expected if retention at the TGN/EE would be affected. However, the acidic residues in the a1-TD could also represent di-acidic ER-export motifs and as we have shown previously that VHA-a3 is targeted to the tonoplast in a Golgi-independent manner (Viotti et al., 2013), we needed to differentiate between the a1-TD serving as an ER exit motif and/or a TGN retention motif. We used inducible expression of the dominant negative GTPase; Sar1BH74L to block ER exit and observed that the mutated VHA-a1 proteins were not retained in the ER and importantly that the signal at the tonoplast was increased. Assuming that the a1-TD contains one or several overlapping diacidic-ER exit motifs and no TGN-retention signal, it should re-route VHA-a3 into COPII- and Golgi-dependent trafficking to the tonoplast. However, a dual localization was observed for VHA-a3-a1-TD indicating either that the a1-TD indeed contains a TGN-retention motif or that VHA-a3 has a yet unknown feature that actively sorts it into the provacuolar route.

### Which VHA-a isoform came first, TGN/EE or tonoplast and are they redundant?

Phylogenetic analysis revealed that the VHA-a isoforms of all angiosperms as well as the gymnosperm *P. taeda* fall into two distinct clades. The a1-TD is conserved within the VHA-a1 clade and exchange of the N-terminal domains within this clade resulted in TGN/EE localization whereas the N-terminal domain of Selaginella that lacks the a1-TD resulted in tonoplast localization indicating that the duplication leading to differential targeting occured in a common ancestor of all seed plants. With the exception of Marchantia, all other plants outside of the spermatophytes possess multiple VHA-a isoforms, however the a1-TD is not conserved in these plants. Evidence for differential targeting of VHA-a isoforms is missing but it seems reasonable to speculate that differential localization arose independently. Based on this, Marchantia might represent the ancestral ground state with a single VHA-isoform predominantly localized at the tonoplast. The trafficking machinery of Marchantia has distinct features and it remains to be determined if acidification of the TGN/EE plays a similar role as in Arabidopsis. If so, Marchantia would indeed represent a unique model for a single VHA-a isoform being sufficient for V-ATPase activity in two locations. A comparable situation has been proposed for the ancestral form of Stv1p and Vph1p that also served both Golgi-and vacuole acidification based on slow anterograde post-Golgi trafficking (Finnigan et al., 2011). In support of this notion, it has recently been shown that V-ATPase complexes containing Vph1p contribute more to Golgi-acidification while en route to the vacuole than the Golgi-localized complexes containing Stv1p (Deschamps et al., 2020). The identity of the VHA-a isoform does not only determine the subcellular localization, it can influence a number of biochemical properties including assembly, coupling efficiency, protein abundance and reversible dissociation (Leng et al., 1998; Kawasaki-Nishi et al., 2001b, 2001a)

Given the central role of the V-ATPase in many cellular functions it is important to not only understand the sorting of this molecular machine but also to address the functional diversification of plant VHA-a isoforms. The fact that VHA-a1 subunits with mutations in the a1-TD which are mislocalized to the tonoplast can complement the dwarf phenotype of the *vha-a2 vha-a3* mutant to varying degrees argues strongly that they have retained their basal proton-pumping activity and do not require the specific lipid-environment of the TGN/EE for functionality. However, VHA-a1 containing V-ATPases at the vacuole cannot acidify the vacuole to wildtype levels and it remains to be determined if this is caused eg. by a difference in coupling efficiency (Kawasaki-Nishi et al., 2001b) or pH-dependent feedback regulation (Rienmüller et al., 2012). In contrast, when the Marchantia VHA-a was expressed in *vha-a2 vha-a3*, the resulting transgenic lines were indistinguishable from wildtype. As null alleles of single copy-encoded VHA-subunits cause male gametophyte lethality (Dettmer et al., 2005), we assumed that we would have to transform heterozygous *vha-a1/+* plants to analyse in their progeny if MpVHA-a and VHA-a3-a1-TD can functionally replace VHA-a1 at the TGN/EE. As we encountered T-DNA related silencing problems in earlier complementation experiments, we used CRISPR/Cas9 under control of an egg cell specific promoter (Wang et al., 2015) to generate new *vha-a1* mutant alleles. Given that we have previously reported that inducible knock-down of *VHA-a1* via either RNAi or amiRNA causes strong cell expansion defects (Brüx et al., 2008), the fact that plants homozygous for several independent null alleles are viable is surprising. Although these individuals were nearly indistinguishable from wildtype during vegetative growth they were found to be completely sterile due to a block in male gametophyte development. Intriguingly, when we systematically reduced the number of wildtype alleles for VHA-a2 and VHA-a3 in the *vha-a1* null background, a vegetative phenotype became manifest indicating that the two tonoplast-isoforms are able to complement for the lack of VHA-a1 during vegetative growth. However, neither VHA-a2, VHA-a3 or MpVHA-a can replace VHA-a1 during pollen development indicating that it has acquired a unique and specialized function in the male gametophyte. To summarize our findings and to address the apparent discrepancy between knock-down and null allele phenotypes, we propose the following model:

### VHA-a1 and VHA-a3 compete for entry into COPII-vesicles

We assume that Marchantia reflects the ancestral state in which ER-export is mediated via COPII and acidification of the TGN/EE is either not required or is provided by V-ATPase complexes passing through en route to the tonoplast. Similar to yeast, the duplication of the ancestral VHA-a isoform provided the basis for a dedicated TGN/EE-isoform which in turn allowed the evolution of a Golgi-independent trafficking route from the ER to the tonoplast, which we have shown previously to exist in meristematic root cells (Viotti et al., 2013). Alternatively, a single VHA-isoform could be able to enter both trafficking routes and it will thus be of importance to determine the evolutionary origin of the provacuolar route.

We propose here that in Arabidopsis, cells that require large quantities of tonoplast membranes such as the cells of the root tip, the VHA-a isoforms compete for entry into COPII-vesicles. Assembly of all V_O_-complexes takes place at the ER with the help of dedicated assembly factors and based on RNA expression and enzymatic activity the ratio of VHA-a1 to VHA-a3 is roughly 1:10 (Hanitzsch et al., 2007; Neubert et al., 2008). We assume that due to the presence of the a1-TD, the affinity of VHA-a1 for Sec24, the cargo receptor for COPII vesicles, is much higher than the affinity of VHA-a3. Small quantities of VHA-a3 can enter COPII vesicles but the majority is predominantly sorted into the Golgi-independent route (Figure 10).The fact that VHA-a3 is strongly trafficked to the TGN/EE by the insertion of the a1-TD provides further evidence for our competition model which also predicts that blocking of COPII-mediated export by DN-Sar1-GTP would lead to re-routing of VHA-a1 to the tonoplast when its affinity for COPII is reduced by point mutations in the a1-TD. The fact that tonoplast-localization of a1-TD mutants is enhanced in the *vha-a2 vha-a3* background provides further support for a competition between the isoforms for the two ER-exits. Similarly, in the absence of VHA-a1, VHA-a2/VHA-a3 would be able to enter the COPII-route more efficiently and would thus be able to compensate for the lack of V-ATPases at the TGN/EE providing an explanation for the lack of a vegetative phenotype of the *vha-a1* mutant. We have shown previously that despite their seemingly strict spatial separation, the activities of the V-ATPase at the TGN/EE and the tonoplast are coordinated. In the *vha-a2 vha-a3* mutant in which VHA-a1 is the only remaining target, treatment with the V-ATPase inhibitor ConcA causes an additional increase in vacuolar pH (Kriegel et al., 2015) although the localization of VHA-a1 is not affected. Why then, does VHA-a3 not enter the COPII route and rescue the reduced activity at the TGN/EE in the RNA-mediated knock-down lines of VHA-a1? Based on our model, we propose that in the RNA-mediated knock-down lines, although levels of VHA-a1 are reduced, they are still sufficient to outcompete VHA-a3 for entry into COPII-vesicles.

**Figure 10:**
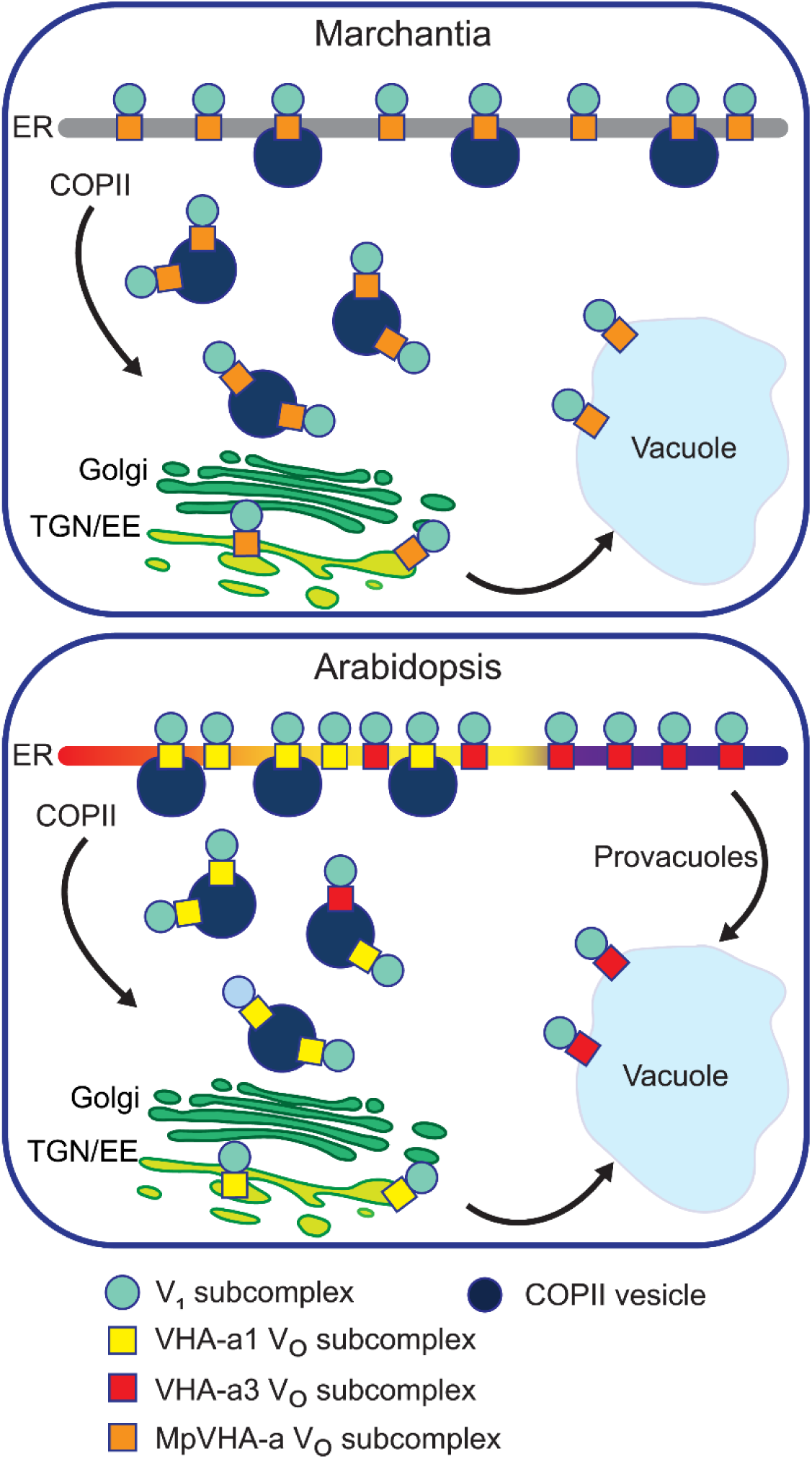
A competition exists to enter COPII vesicles in Arabidopsis. Upper panel: COPII-mediated ER-export is the only way to exit the ER in Marchantia. Marchantia VHA-a (MpVHA-a) does not contain a TGN/EE retention signal. MpVHA-a complexes acidify the TGN/EE en route to the tonoplast. Lower panel: Two ER exits exist in Arabidopsis; COPII-mediated ER-export and exit via provacuoles. A competition exists between VHA-a1 and VHA-a3/VHA-a2 complexes to enter COPII vesicles. VHA-a1 has a higher affinity for COPII machinery conferred by the a1-TD. Therefore, it is preferentially loaded into COPII vesicles over VHA-a3. VHA-a3 containing complexes are transported to the tonoplast via provacuoles.

In conclusion, our study provides the first report on a targeting motif for subunit a from a multicellular eukaryotic V-ATPase. Interaction partners that recognize and bind to the a1-TD need to be identified. ER exit motifs as well as provacuole directing motifs in VHA-a3 should also be uncovered to better understand its trafficking via COPII vesicles and provacuoles. Our results incite further investigations to determine additional functions of the endosomal V-ATPase during vegetative growth and to learn what new functions it acquired during evolution that are essential for pollen development.

## Experimental procedures

### Plant materials and growth conditions

*Arabidopsis thaliana*, Columbia 0 (Col-0) ecotype was used in all experiments in this study. The *vha-a2 vha-a3* double mutant was characterized by Krebs et al., 2010. *VHA-a1p:VHA-a1-GFP*, *VHA-a3p:VHA-a3-GFP* and *VHA-a1p:VHA-a1-RFP* lines were established by Dettmer et al., 2006. *BRI1-GFP* lines were previously established by Geldner et al., 2007.

Growth of Arabidopsis seedlings for confocal microscopy was performed on plates. The standard growth medium used contained 1/2 Murashige and Skoog (MS), 0.5 % sucrose, 0.5 % phyto agar, 10 mM MES and the pH was set to 5.8 using KOH. Agar and MS basal salt mixture were purchased from Duchefa. Seeds were surface sterilized with ethanol and stratified for 48h at 4°C. Plants were grown in long day conditions (LD; 16 h light/8 h dark) for 5 days.

For the rosette phenotype assays, seeds were stratified for 48 h at 4°C and then placed on soil. Seedlings were transferred to individual pots at 7 days after germination (DAG). Plants were grown either in LD or short day conditions (SD; 10 h light/14 h dark) for the required time.

*M*. *polymorpha* accession Takaragaike-1 (Tak-1, male; Ishizaki et al., 2008) was used in this study. The growth condition and the transformation method were described previously (Kubota et al., 2013; Kanazawa et al., 2016). *M*. *polymorpha* expressing mRFP-MpSYP6A was generated in the previous work (Kanazawa et al., 2016).

### Construct design and plant transformation

#### VHA-a1/VHA-a3 Chimeras

Five chimeric proteins were made which consisted of increasing lengths of the VHA-a1 N-terminus (37aa, 85aa, 131aa, 179 aa and 228 aa) fused to decreasing lengths of the C-terminal domain of VHA-a3. The GreenGate cloning system was used (Lampropoulos et al., 2013). Unique overhangs for each chimera were designed to allow seamless fusion of the *VHA-a1* and *VHA-a3* cDNA sequences (Supplemental Table 4). The primers used are listed in Supplemental Table 4. All PCR products were blunt end cloned into the pJET1.2 vector (ThermoFisher Scientific) and verified by sequencing (Eurofins). Verified clones were digested using either *Bgl*ll *or Not*I *and Cla*I to release the fragments. To combine the *VHA-a1* and *VHA-a3* fragments, the GreenGate cloning system was applied using modules described in Supplemental Table 5.

#### UBQ:VHA-a3-a1-TD-GFP

The targeting domain of *VHA-a1* (*a1-TD*) was introduced into *VHA-a3* by PCR techniques. The N and C-termini of *VHA-a3* and the *a1-TD* were amplified using primers in Supplemental Table 4. All PCR products were blunt end cloned into the pJET1.2 vector and verified by sequencing (Eurofins). Verified clones were digested using either *Bgl*ll *or Not*I *and Cla*I to release the fragments. The destination vector was made by combining the fragments with GreenGate modules described in Supplemental Table 5.

#### Site-directed mutagenesis of VHA-a1

The pJET1.2 clone carrying VHA-a1 NT with 179 aa from the chimeras was used as a template for site directed mutagenesis of VHA-a1. PCR mediated site-directed mutagenesis was performed using primers indicated in Supplemental Table 4 and according to the manufacturer’s protocol. Arabidopsis *VHA-a1 intron 10* was amplified from genomic DNA, a connecting *VHA-a1* fragment (*Y2*) and a C-terminal fragment (*VHA-a1-CT*) were amplified from *VHA-a1* cDNA using primers indicated in Supplemental Table 4. All PCR fragments were subcloned into the pJet1.2 vector and were verified by sequencing (Eurofins). Verified clones were digested using *Bgl*ll to release the fragments. The final destination vectors were made with modules described in Supplemental Table 5.

#### UBQ:VHA-a3R729N-GFP

A 364 bp fragment was excised from the VHA-a3 cDNA with *SalI* and *EcoRI* and subcloned into the pJet1.2 vector generating the plasmid pa3Cterm. Site directed mutagenesis was performed on the pa3Cterm according to the manufacturer’s protocol using primers R729N_for and R729N_rev (Supplemental Table 4). The mutation was verified by sequencing (Eurofins). Verified clones were released using *Sal*I and *Eco*RI restriction sites and inserted back into the pUGT2kan containing the wildtype *VHA-a3* cDNA.

#### Dex:Sar1BH74L-CFP

The Arabidopsis *Sar1B* cDNA sequence with the H74L mutation was synthesized by Eurofins. The synthesized fragment contained *Eco*31I at its 5’ and 3’ ends for subcloning. PCR was performed on the synthesized fragment using primers indicated in Supplemental Table 4 and the PCR product was blunt end cloned into the pJET1.2 vector. The *SarB1H74L* sequence was verified by sequencing (Eurofins). Verified clones were digested using *Not*I *and Cla*I to release the *SarB1H74L* fragments. The dexamethasone inducible construct was made by using the LhG4/pOp system combined with the ligand-binding domain of the rat glucocorticoid receptor (GR; Moore et al., 1998; Craft et al., 2005; Samalova et al., 2005). Two GreenGate reactions were performed to create two intermediate vectors that were later combined on one T-DNA. The first intermediate vector (pKSM002) contained the GR-LhG4 transcription factor expressed under the *UBQ10* promoter and the second intermediate vector contained *Sar1BH74L* under the *pOP6* promoter (pKSN009). The two intermediate vectors were combined on one final destination vector pGGZ003 (Supplemental Table 5).

#### UBQ10:MpVHA-a-mVenus

The cDNA sequence of *MpVHA-a* was amplified by PCR using female *Marchantia polymorpha* (ecotype BoGa) thallus cDNA as a template. The sequence was amplified in two parts (*MpVHA-aNT and MpVHA-aCT)* separated at the exon 10-exon 11 junction using primers indicated in Supplemental Table 4. All PCR products were blunt end cloned into the pJET1.2 vector and verified by sequencing (Eurofins). Verified clones were digested using *Bgl*ll to release the fragments. The final destination vector was made with the *MpVHA-aNT, VHA-a1-intron10 and MpVHA-aCT* fragments and modules described in Supplemental Table 5.

#### CaMV35S:MpVHA-a-mVenus and MpEF1α:MpVHA-a-mVenus

The genome sequence of *MpVHA-a* containing all exons and introns was amplified by PCR using the Tak-1 genome as a template with primers (Supplemental Table 4). The amplified products were subcloned into pENTR D-TOPO (Invitrogen), and cDNA for *mVenus* was inserted into the *Asc*I site of the pENTR vectors using In-Fusion (Clontech) according to the manufacturer’s instructions. The resultant entry sequences were transferred to pMpGWB302 or pMpGWB303 (Ishizaki et al., 2015) using LR Clonase II (Invitrogen) according to the manufacturer’s instructions.

#### UBQ10:A.trichopoda-VHA-aNT-VHA-a1-mCherry,UBQ10:S.moellendorffii-VHA-aNT-VHA-a1-mCherry and UBQ10:P.taeda-VHA-aNT-VHA-a1-mVenus

The first 682 bp of the sequence identified as evm_27.model.AmTr_v1.0_scaffold00080.37 (*A. trichopoda VHA-a*), 649 bp of the sequence identified as 182335 (*S.moellendorffii-VHA-a*) and a 703 bp fragment starting from the 163rd bp of the sequence identified as 5A_I15_VO_L_1_T_29156/41278 (*P.taeda-VHA-a*) were synthesized (Eurofins). The synthesized fragments contained *Eco*31I at their 5’ and 3’ ends for subcloning. PCR was performed on the synthesized fragments using primers indicated in Supplemental Table 4 and the PCR products were blunt end cloned into the pJET1.2 vector. A connecting *VHA-a1* fragment (dodo) was also amplified from *VHA-a1* cDNA using primers in Supplemental Table 4. All PCR fragments were verified by sequencing (Eurofins). Verified clones were digested using either *Bgl*ll or *Not*I *and Cla*I to release the fragments. The destination vectors were made by combining the following components: *A.trichopoda-VHA-aNT* or *P.taeda-VHA-aNT* or *S.moellendorffii-VHA-aNT* with *VHA-a1 dodo*, *VHA-a1-intron10*, *VHA-a1-CT* and modules described in Supplemental Table 5.

#### UBQ10:ST-GFP

Rat *Sialyltransferase* (ST) sequence was amplified from pre-existing p16:ST-pHusion plasmid (Luo et al., 2015) using GG-ST-C-fwd and GG-ST-C-rev primers (Supplemental Table 4), sub-cloned into pGGC000 after *Eco*31I digest and sequenced to verify correct insert sequence. pGGC-ST was used in a GreenGate reaction to generate *UBQ10:ST-GFP* using modules described in Supplemental Table 5.

#### UBQ10:VHA-a3-pmScarlet-I

The *VHA-a3* cDNA sequence was amplified from cDNA with primers listed in Supplemental Table 4. After digest with *Eco*31I, VHA-a3 was subcloned into pGGC000. pGGC-VHA-a3 was used in a GreenGate reaction to assemble the final destination vector as described in Supplemental Table 5.

#### UBQ10:VHA-a1-GFP

The *VHA-a1* cDNA sequence was split into two parts by PCR to generate the fragments *VHA-a1 NT (Y5)* and *VHA-a1 CT* separated at the exon 10-exon 11 junction using primers indicated in Supplemental Table 4. These fragments were digested with *Eco*31I and then subcloned into pGGC000 with *VHA-a1 intron 10*. pGGC-VHA-a1-intron 10 was sequenced and used in a GreenGate reaction with modules described in Supplemental Table 5 to generate the final destination vector.

#### Destination vector pGGZ004

The vector backbone pGGZ004 is a GreenGate compatible plant binary vector which, unlike previous GreenGate vector backbones (Lampropoulos et al., 2013), contains the replicase required for plasmid replication in *A. tumefaciens* and therefore does not require the pSOUP helper plasmid (Hellens et al., 2000). pGGZ004 is based on the plant binary vector pTKan (Krebs et al., 2012), which originated from pPZP212 (Hajdukiewicz et al., 1994). To generate pGGZ004, pTKan was cut at its multiple cloning site with *Kpn*I followed by an incomplete digest with *Eco31*I, which opened the plasmid next to the T-DNA left border. The resulting 6971-bp fragment comprised the T-DNA border regions, the pBR322 *bom* site and the ColE1 and pVS1 plasmid origins for replication in *E. coli* and in *Agrobacterium*, respectively (Hajdukiewicz et al., 1994). GreenGate overhangs A and G were added to the opened vector backbone by sticky end ligation of annealed oligonucleotides *Eco31*I-w-AG-Fw and *Eco31*I-w-AG-Rv which contained *Kpn*I and *Eco31*I compatible overhangs. Lastly, a site-directed mutagenesis using primer combination pGGZ004SDM1-Fw and pGGZ004SDM1-Rv was performed to remove an *Eco31*I site in the vector backbone (Supplemental Table 4). The final pGGZ004 vector was fully sequenced before use for further cloning.

#### CRISPR/Cas9 constructs

CRISPR target sites in *VHA-a1* were selected using CHOPCHOP (https://chopchop.cbu.uib.no; Labun et al., 2016) and CCtop; https://crispr.cos.uni-heidelberg.de, Stemmer et al., 2015). gRNAs with at least 4 bp difference to every other region in the *Arabidopsis thaliana* genome were selected as precaution against off-target mutations. gRNA sequences (Supplemental Table 6) were inserted into the plasmid pHEE401E as described by the authors (Wang et al., 2015). Three CRISPR plasmids were cloned, one containing two gRNAs (gRNA1 and gRNA2), aiming at deleting the region between the two CRISPR sites, and two plasmids containing one gRNA each, gRNA3 and gRNA4.

Depending on the final destination vector used, all constructs were transformed into either of two *Agrobacterium tumefaciens* strains. The strain GV3101:pMP90 was used if the final destination vector was pGGZ004. Selection was done on 5 mg/ml rifampicin, 10 mg/ml gentamycin, and 100 mg/ml spectinomycin. The strain ASE1(pSOUP+) was used if pGGZ001/3 were used. Selection was done on 100 μg/ml spectinomycin, 5 μg/ml tetracycline (for pSOUP), 25 μg/ml chloramphenicol and 50 μg/ml kanamycin. Arabidopsis plants were transformed using standard procedures. Transgenic plants were selected on MS medium containing appropriate antibiotics.

### Analysis of mutations at CRISPR sites and genotyping of mutants

Genomic DNA was extracted from rosette leaves and amplified by PCR using primers flanking the CRISPR sites (Supplemental Table 6). PCR products were sequenced by Eurofins or analyzed by agarose gel electrophoresis. Presence/absence of CRISPR T-DNA was monitored by PCR with *Cas9* specific primers (Supplemental Table 6) followed by agarose gel electrophoresis. *VHA-a2* and *VHA-a3* wildtype and mutant alleles were identified by PCR using specific primers (Supplemental Table 6) and agarose gel electrophoresis. Screening for *vha-a1-1* was done by detecting the PCR product spanning the CRISPR sites (260 bp shorter for *vha-a1-1* compared to wildtype) by agarose gel electrophoresis (Supplemental Figure 12B). Screening for *vha-a1* mutants with other *vha-a1* alleles was performed by observation of the pollen phenotype after having established that the defect in pollen development is caused by knockout of *VHA-a1* (Supplemental Table 2).

### Pharmacological Treatments and Stains

Arabidopsis seedlings were incubated in liquid 1/2 MS medium with 0.5 % sucrose, pH 5.8 (KOH), containing 50 µM BFA, 1 µM FM4-64, 60 µM DEX and 125 nM ConcA or the equivalent amount of DMSO in control samples for the required time at room temperature. Stock solutions were prepared in DMSO.

Alexander’s stain: Anthers from flowers at stage 12 (Smyth et al., 1990) were incubated in Alexander’s stain (Alexander, 1969) on objective slides covered with coverslips at room temperature for 20 hours. Bright field images were taken using a Zeiss Axio Imager M1 microscope.

### Confocal Microscopy

Arabidopsis root cells of 6day-old seedlings were analyzed by confocal laser scanning microscopy (CLSM) using a Leica TCS SP5II microscope equipped with a Leica HCX PL APO lambda blue 63.0x 1.20 UV water immersion objective. CFP, GFP and mVenus were excited at 458 nm, 488 nm and 514 nm with a VIS-argon laser respectively. mRFP, mCherry, pmScarlet-I and FM4-64 were excited at 561 nm with a VIS-DPSS 561 laser diode. For image acquisition, the Leica Application Suite Advanced Fluorescence software was used. Processing of images was performed using the Leica LAS AF software.Gaussian blur with kernel size 3 was applied. Average tonoplast intensities were measured in roots of Arabidopsis seedlings expressing *DEX:Sar1BH74L-CFP* and *UBQ10:VHA-a1-GFP* with mutations or *VHA-a3:VHA-a3-GFP* after 6 hours induction with 60 µM DEX (Sigma-Aldrich). Images were acquired sequentially and with identical settings. In the first sequential scan, GFP was excited at 488 nm and emission detected at 500-545 nm. In the second sequential scan CFP was exited at 458 nm and emission detected between 470-485 nm. Intensity measurements were done using plot profiles in ImageJ. The maximum intensity values along line profiles across the TGN/EE and tonoplast were recorded. OriginPro was used to determine if the data were normally distributed and to perform statistical tests. To determine the ratio of TGN/EE-to-tonoplast fluorescence intensity, images of root cells expressing UBQ10:VHA-a1-GFP with mutations in the wildtype and vha-a2 vha-a3 background were acquired using identical settings. The maximum intensity values along line profiles across the TGN/EE and tonoplast were measured using ImageJ. The average TGN/EE and tonoplast intensities were calculated for each image and the TGN/EE-to-tonoplast fluorescence intensity ratio was calculated. The TGN/EE-to-tonoplast fluorescence intensity ratios for n ≥10 images for each mutation were averaged and statistical tests were performed with OriginPro. CLSM on Marchantia thalli was performed according to (Kanazawa et al., 2016). Briefly, the dorsal cells of 5-day-old thalli were observed using LSM780 (Carl Zeiss). Spectral linear unmixing of obtained images was performed using ZEN2012 software (Carl Zeiss).

### High-Pressure Freezing, Freeze Substitution, and EM

Seven-day-old Arabidopsis wildtype seedlings expressing *DEX:AtSar1b-GTP* were induced for 6 hours with 60 μM DEX. Seedlings were then processed as previously described (Scheuring et al., 2011). Freeze substitution was performed in a Leica EM AFS2 freeze substitution unit in dry acetone supplemented with 0.3% uranyl acetate as previously described (Hillmer et al., 2012). Root tips were cut axially with a Leica Ultracut S microtome to obtain ultrathin sections. Sections were examined in a JEM1400 transmission electron microscope (JEOL) operating at 80 kV. Micrographs were recorded with a TemCam F416 digital camera (TVIPS, Gauting, Germany).

### pH Measurements

Cell sap pH measurements were performed as previously described (Krebs et al., 2010).

### Tonoplast membrane preparation

Tonoplast membranes were prepared from rosette leaves of 6-week-old plants grown under short day conditions as previously described by (Barkla et al., 1999; Leidi et al., 2010).

### SDS-PAGE and Immunoblotting

Microsomal membrane and tonoplast membrane proteins were analyzed by SDS-PAGE and subsequent immunoblotting. Upon gel electrophoresis, the proteins were transferred to a PVDF membrane (Bio-Rad). The primary antibodies against VHA-a1 (AS142822) and VHA-a3 (AS204369) were purchased from Agrisera and were used in a dilution of 1:1000 in 2% BSA-TBS-T. The primary antibody against VHA-B (Ward et al., 1992) and anti-GFP (Roth et al., 2018) were previously described. Antigen on the membrane was visualized with horseradish peroxidase-coupled anti-rabbit IgG (Promega) for VHA-a1 and VHA-a3, anti-mouse IgG (Sigma) for VHA-B and chemiluminescent substrate (Peqlab). Immunostained bands were analyzed using a cooled CCD camera system (Intas).

### Phylogenetic analysis

For phylogenetic reconstruction in a first step the best molecular evolutionary model was determined by running the program PartitionFinder 2 (Lanfear et al., 2017). Phylogenetic reconstruction was then performed by running raxml-ng (Kozlov et al., 2019) setting the model to JTT+I+G and starting the analysis from 10 most parsimonious and 10 random trees. In order to estimate the reliability of the phylogenetic reconstruction 500 bootstrap replicates were run.

### Multiple sequence alignments

Multiple sequence alignments were performed using Clustal omega (Madeira et al., 2019). Aligned sequences were analyzed in Geneious 10.1.3.

### Homology modelling

3D models of the VHA-a1 and VHA-a3 N-termini were obtained through homology modelling with cryo-EM derived models of Stv1p-V_O_ subcomplex (PDB6O7U) and Vph1p-V_O_ subcomplex (PDB6O7T) as templates (Vasanthakumar et al., 2019). Homology modelling was performed according to Roy et al., 2010.

### Accession Numbers

Sequence data from this article can be found in the Arabidopsis Genome Initiative or GenBank/EMBL databases under the following accession numbers: VHA-a1, At2g28520; VHA-a3, At4g39080; Sar1B, AT1G56330.1. Data for *Marchantia polymorpha* VHA-a can be found on the Marchantia genome database with the following ID: Mp3g15140.1. *Amborella trichopoda* and *Selaginella moellendorffii* VHA-a sequences can be found on the Phytozome platform (Goodstein et al., 2012) with the following identifiers: *A.trichopoda* VHA-a; evm_27.TU.AmTr_v1.0_scaffold00080.37 and *S.moellendorffii* VHA-a; 182335. *Pinus taeda* sequence data can be found on the PineRefSeq project on the TreeGenes platform (Falk et al., 2019; Wegrzyn et al., 2008) with the following identifier: 5A_I15_VO_L_1_T_29156/41278.

## Acknowledgements

We would like to thank Prof. Dr. Sabine Zachgo (University of Osnabrück) for kindly providing us with *Marchantia polymorpha* (ecotype BoGa) cDNA. We also thank Prof. Dr. Stephan Rensing (University of Marburg) for providing us with *Selaginella moellendorffii* leaf tissue. We thank Dr. Rainer Waadt (COS, University of Heidelberg) for kindly providing GreenGate entry vectors. We thank Dr. Takayuki Kohchi, Dr. Ryuichi Nishihama (Kyoto University), and Dr. Kimitsune Ishizaki (Kobe University) for vectors. We also thank Dr. Kazuo Ebine, Mayuko Yamamoto, Koji Hayashi (NIBB) and Fabian Fink (COS, University of Heidelberg) for supporting experiments. The support in *M*. *polymorpha* cultivation was provided by the Model Plant Research Facility of National Institute for Basic Biology. This work was supported by the Deutsche Forschungsgemeinschaft within SFB1101 to K.S and by Grants-in-Aid for Scientific Research from the Ministry of Education, Culture, Sports, Science, and Technology of Japan (to T.U., 19H05675, and 18H02470, and T.K., 18K14738). Electron microscopy was performed by S. Hillmer at the Electron Microscopy Core Facility of Heidelberg University with the technical assistance of S. Gold.

## Author Contributions

U.L., R.R., J.A. and K.S. designed experiments. U.L., R.R., J.A. performed experiments. U.L. and M.K. analyzed data. C.K. performed the phylogenetic analysis. T.K. carried out the experiments in Marchantia. S.H. performed electron microscopy. T.U., T.K. and M.K. made comments on the manuscript. U.L., R.R. and K.S. wrote the manuscript.

## Supplemental data

**Supplemental Figure 1.**
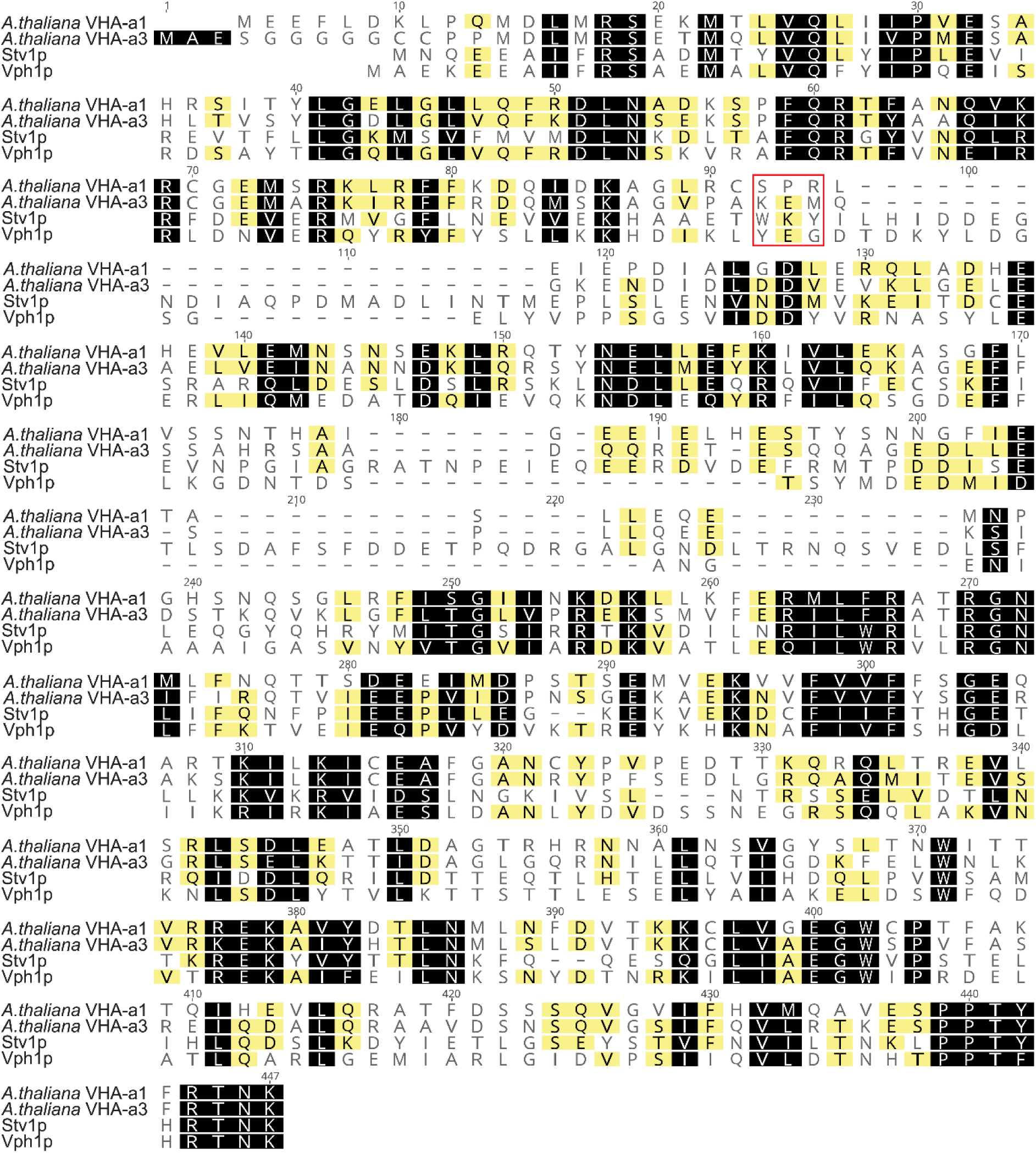
The tri-peptide motif that is responsible for the targeting of Stv1p is absent in VHA-a1. Amino acid sequence alignment of the yeast subunit a isoforms Vph1p and Stv1p with the Arabidopsis isoforms VHA-a1 and VHA-a3. Residues similar between all proteins at the same position are shown against a black background. Residues similar between only three of the proteins at the same position are shown against a grey background. Un-highlighted residues have no similarity in all four sequences at the same position. The position of the tri-peptide motif that is responsible for the targeting of Stv1p is indicated with a red box.

**Supplemental Figure 2.**
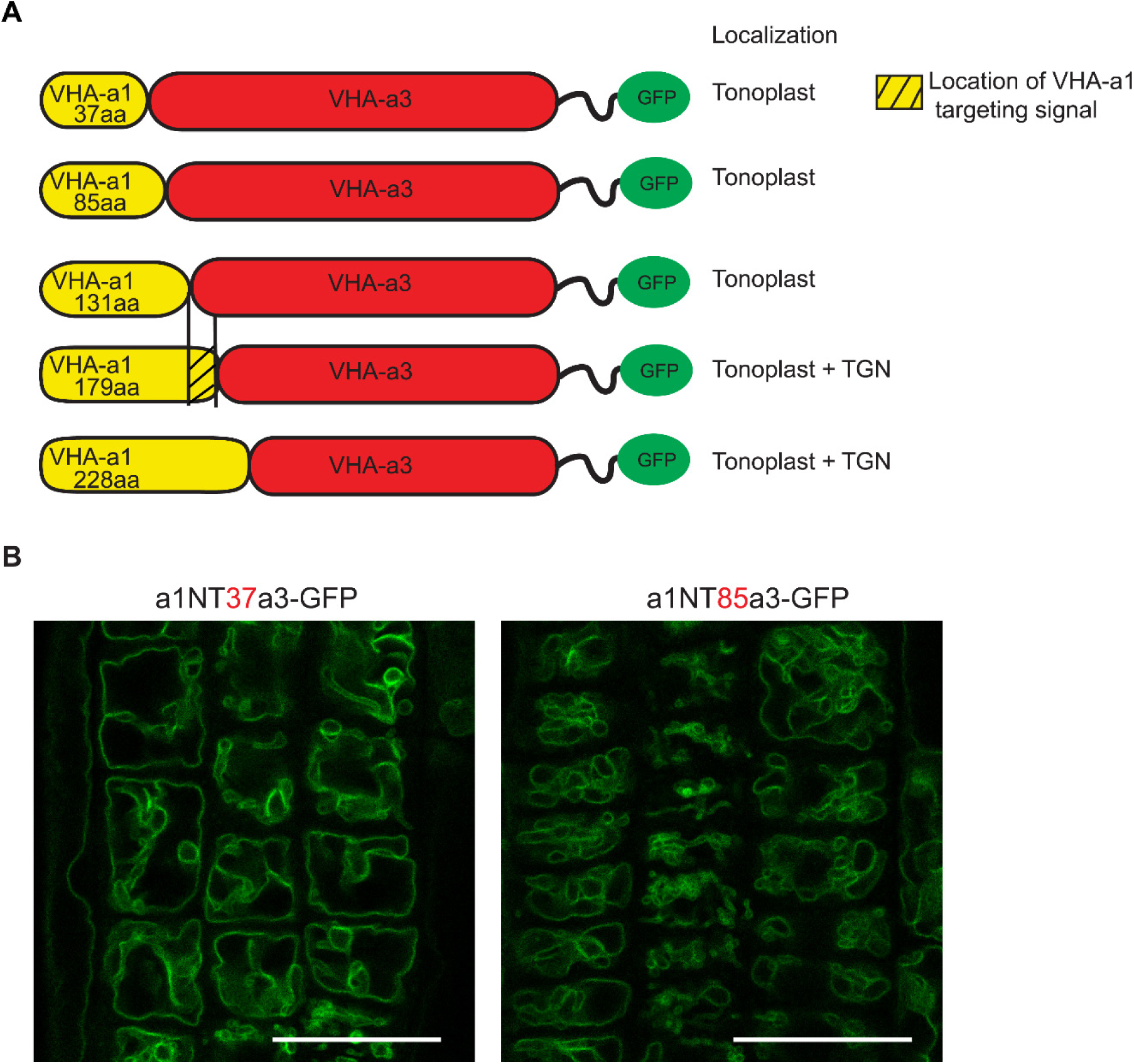
The targeting signal of VHA-a1 is not located in the first 85 amino acids. Chimeric proteins were made which consisted of increasing lengths of the VHA-a1 N-terminus fused to decreasing lengths of the C-terminal domain of VHA-a3. All constructs were fused to GFP. **(A)** Root tips of 6-day-old seedlings were analyzed CLSM. The first two chimeric constructs consisting of 37 aa and 85 aa of the VHA-a1 N-terminal domain localized at the tonoplast. Scale bars = 25 µm.

**Supplemental Figure 3.**
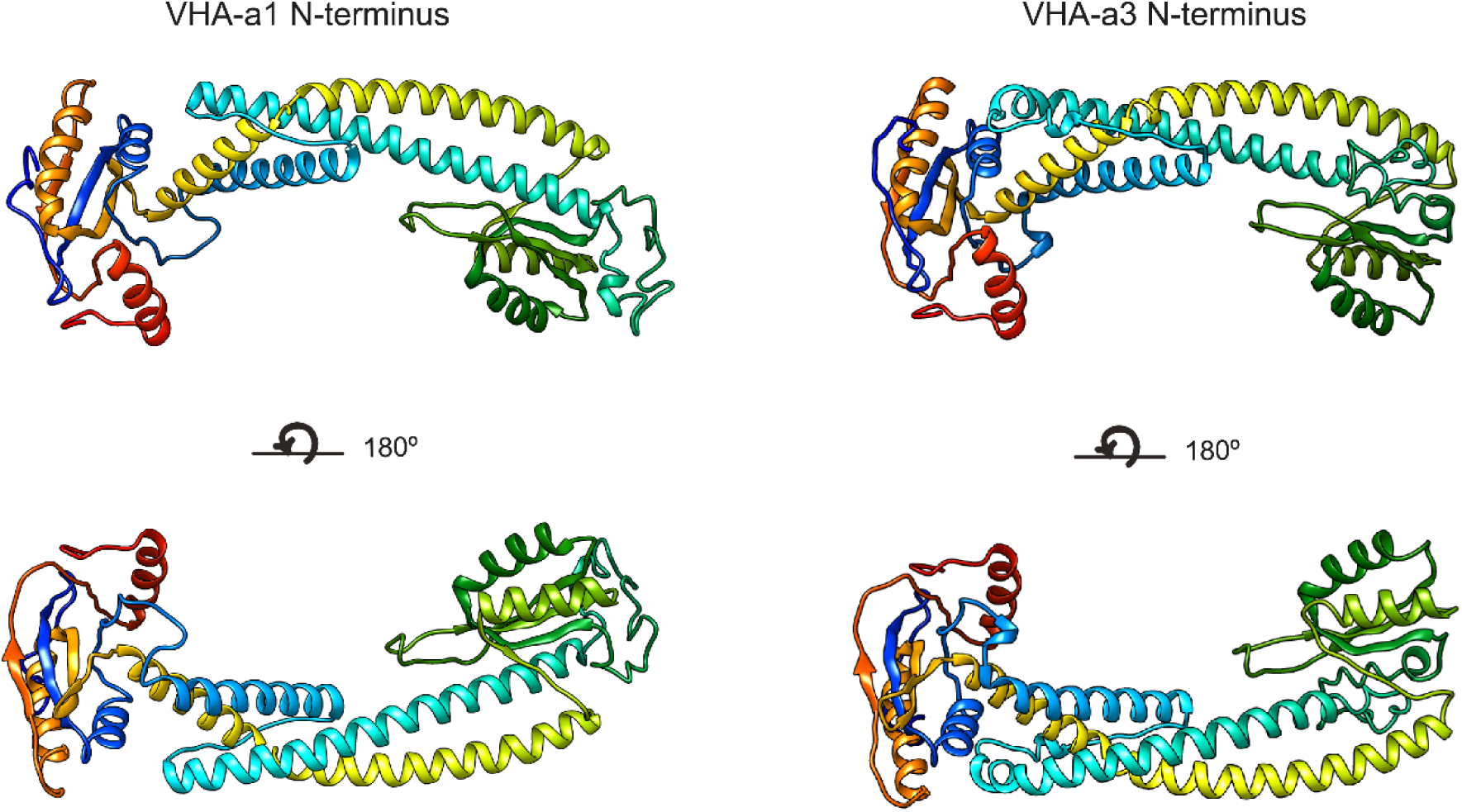
Three-dimensional models of the VHA-a1 and VHA-a3 N termini. Homology modelling of the N-termini of VHA-a1 and VHA-a3 was done using cryo-EM models of Stv1p (PDB607U) and Vph1p (PDB607T) respectively as templates. The models are colour ramped.

**Supplemental Figure 4.**
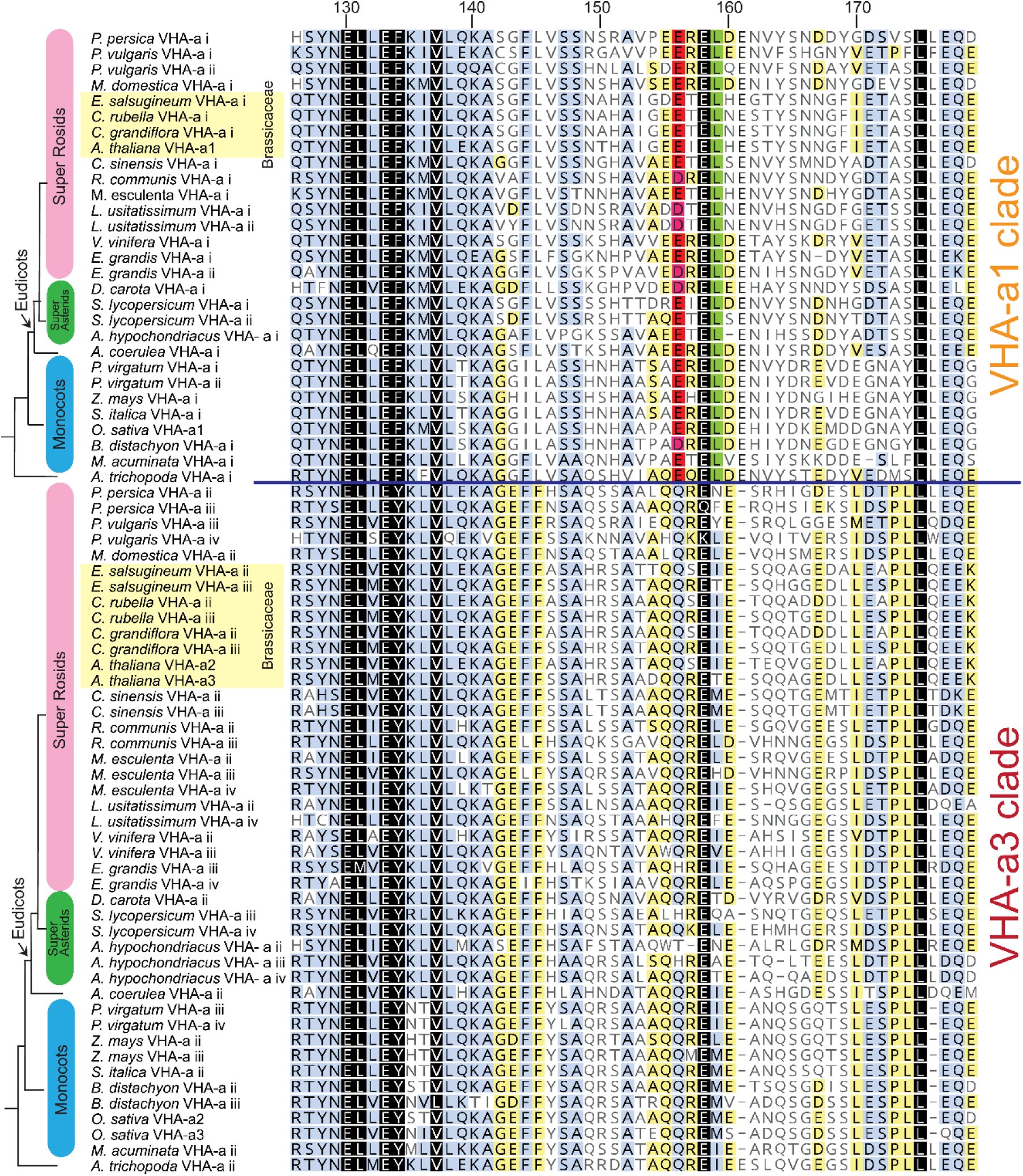
The VHA-a1-TD is conserved in Angiosperms. Amino acid sequence alignment of representative sequences from the VHA-a1 and VHA-a3 clades. The sequence numbers are in reference to the *A. thaliana* VHA-a1 sequence. Residues highlighted in black, blue and yellow are 100%, 80-100% and 60-80% similar respectively. Un-highlighted residues are less than 60% similar. The alignment is restricted to the VHA-a1 targeting domain (His126 to Glu 179 in *A. thaliana* VHA-a1). The alignment reveals that there is an acidic cluster that is only present in the VHA-a1 clade.

**Supplemental Figure 5.**
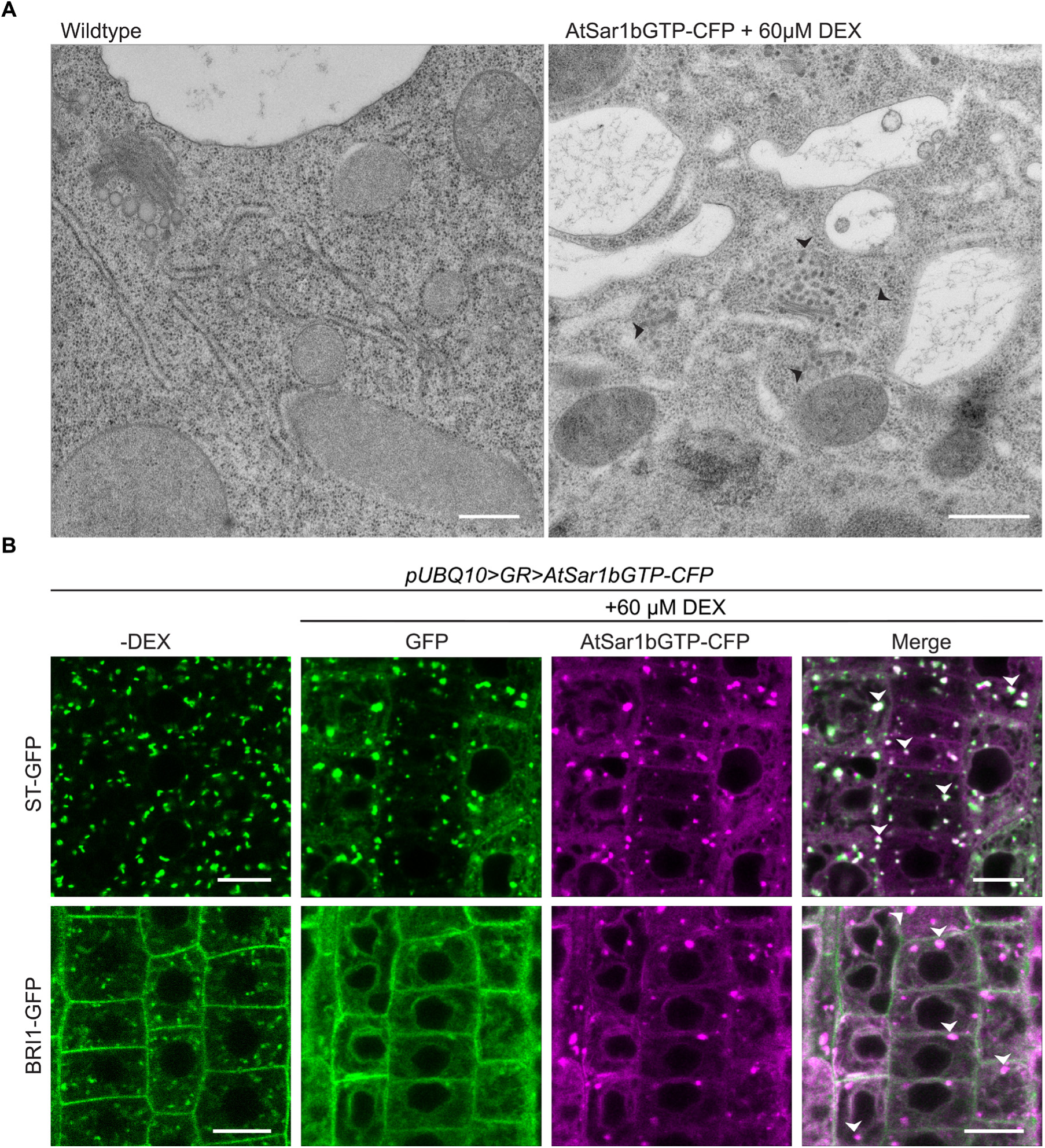
AtSar1b-GTP-CFP expression blocks the ER exit of secretory pathway proteins. **(A)** EM of high-pressure frozen root tips. AtSar1b-GTP-CFP expression causes bloating of the ER, aggregation of Golgi stacks and leads to an accumulation of vesicles in the cell (black arrows). Scale bars = 500 nm. **(B)** After 6 hours of induction with 60 µM DEX, AtSar1b-GTP-CFP is expressed. Both the Golgi targeted protein; ST-GFP and plasma membrane destined BRI1-GFP are retained at the ER when exit from the ER via COPII vesicles is blocked by expression of AtSar1b-GTP-CFP. Scale bars = 10 µm.

**Supplemental Figure 6.**
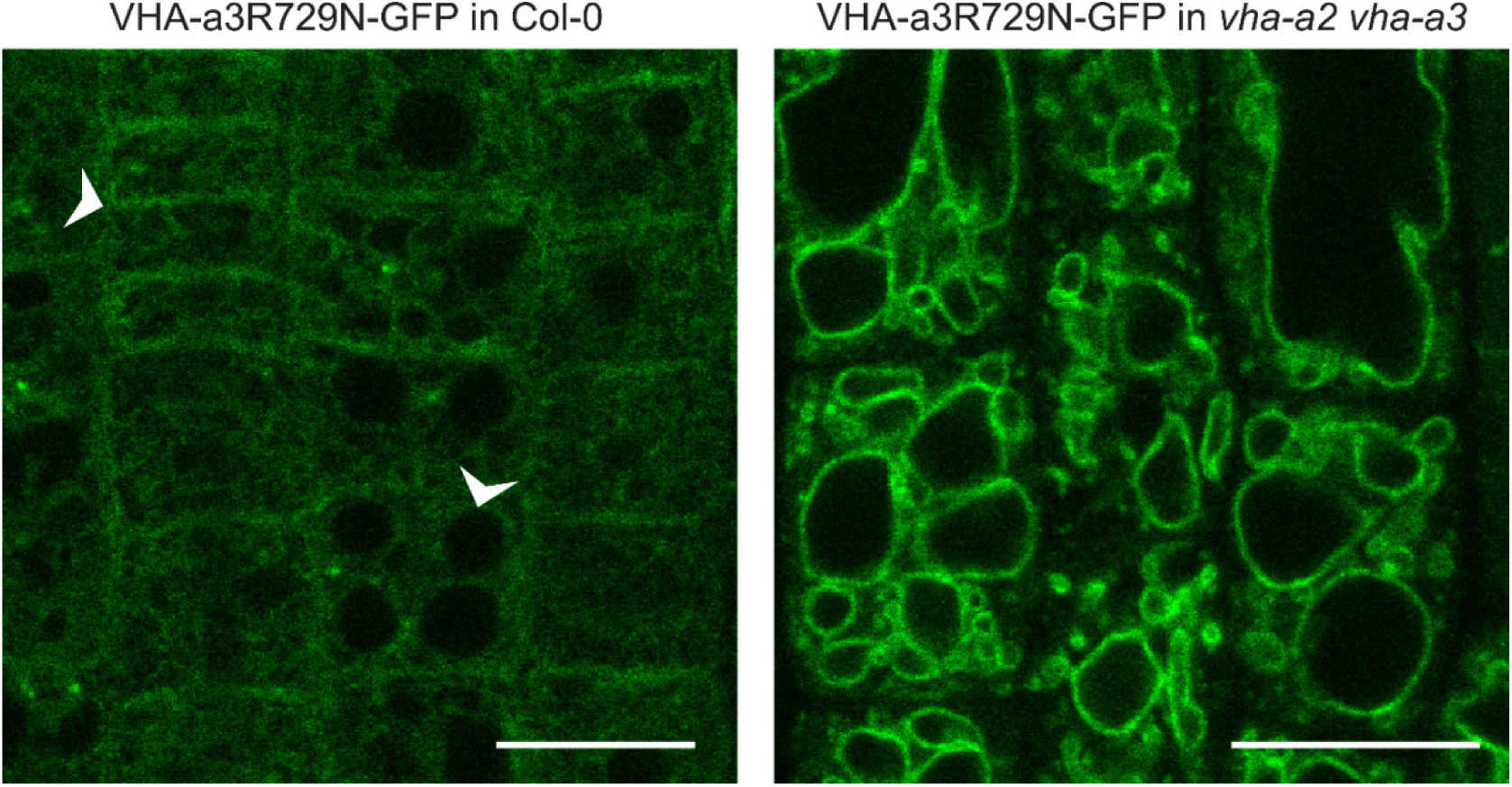
A competition exists to enter provacuoles in wildtype *Arabidopsis* root tip cells. VHA-a3-R729N-GFP is retained in the wildtype background and localizes to the tonoplast in the *vha-a2 vha-a3* double mutant background.

**Supplemental Figure 7.**
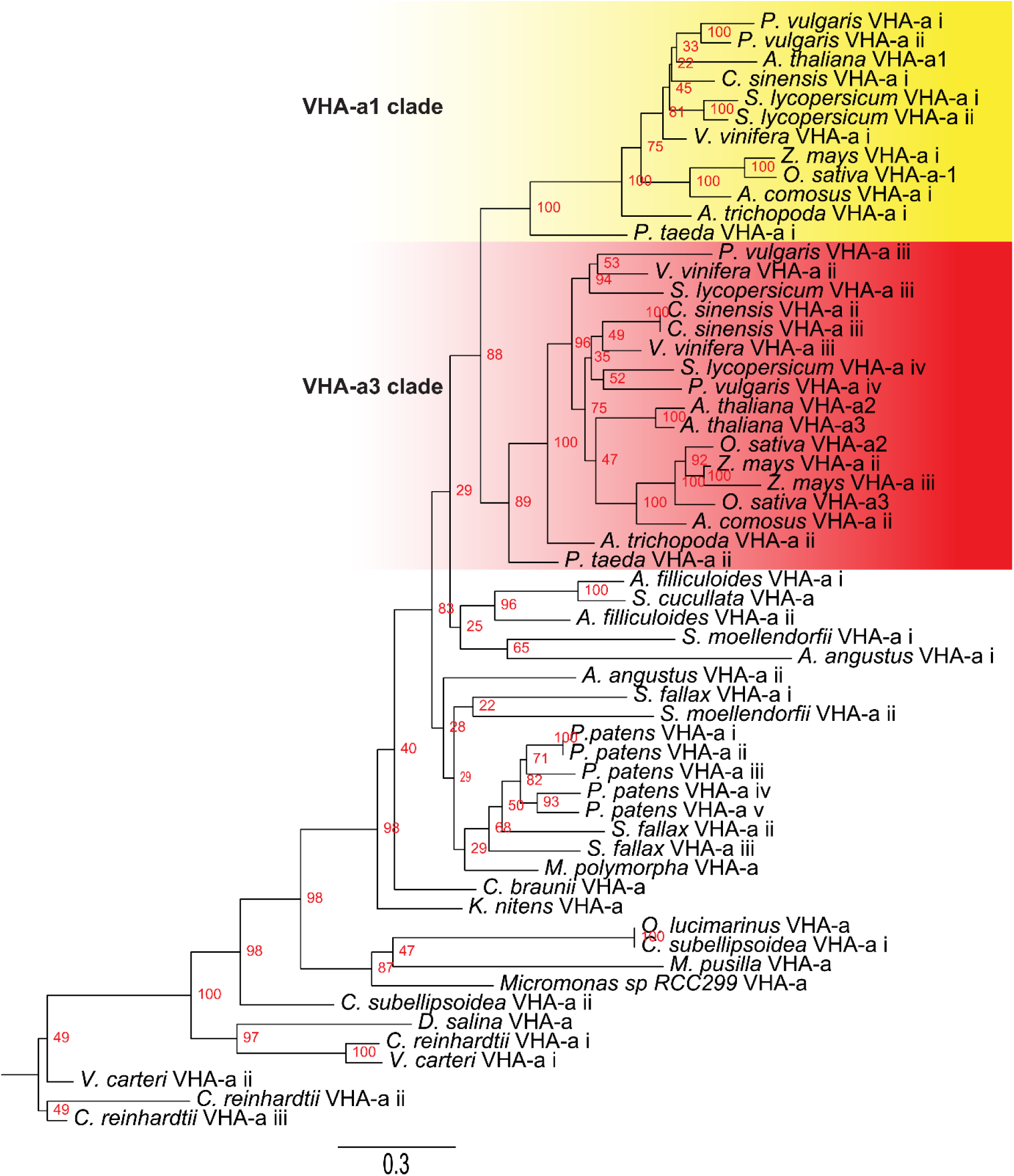
Phylogenetic analysis of the N-terminal sequences of VHA-a related proteins. VHA-a related protein sequences for selected species are shown. Branch support is calculated on the basis of 500 bootstraps.

**Supplemental Figure 8.**
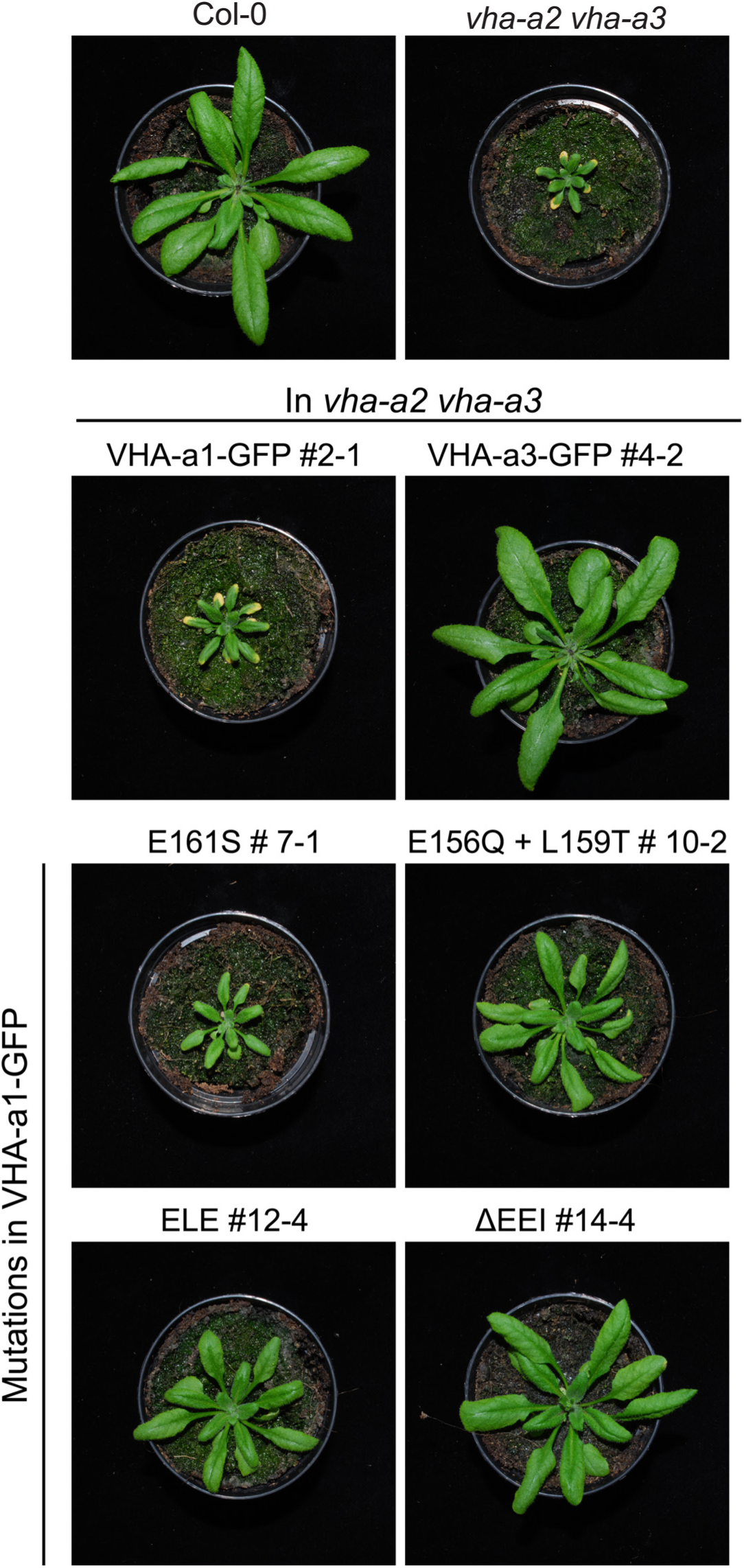
The mutated VHA-a1-GFP proteins complement the *vha-a2 vha-a3* double mutant to varying degrees in long day conditions. Plants were grown in long day conditions (22°C and 16 hours light) for 4 weeks. All mutant variants of VHA-a1-GFP displayed bigger rosette size than the *vha-a2 vha-a3* double mutant. VHA-a1 with the ΔEE1 mutation (E155+ E156 + I157 deletion) complements the *vha-a2 vha-a3* double mutant the best.

**Supplemental Figure 9.**
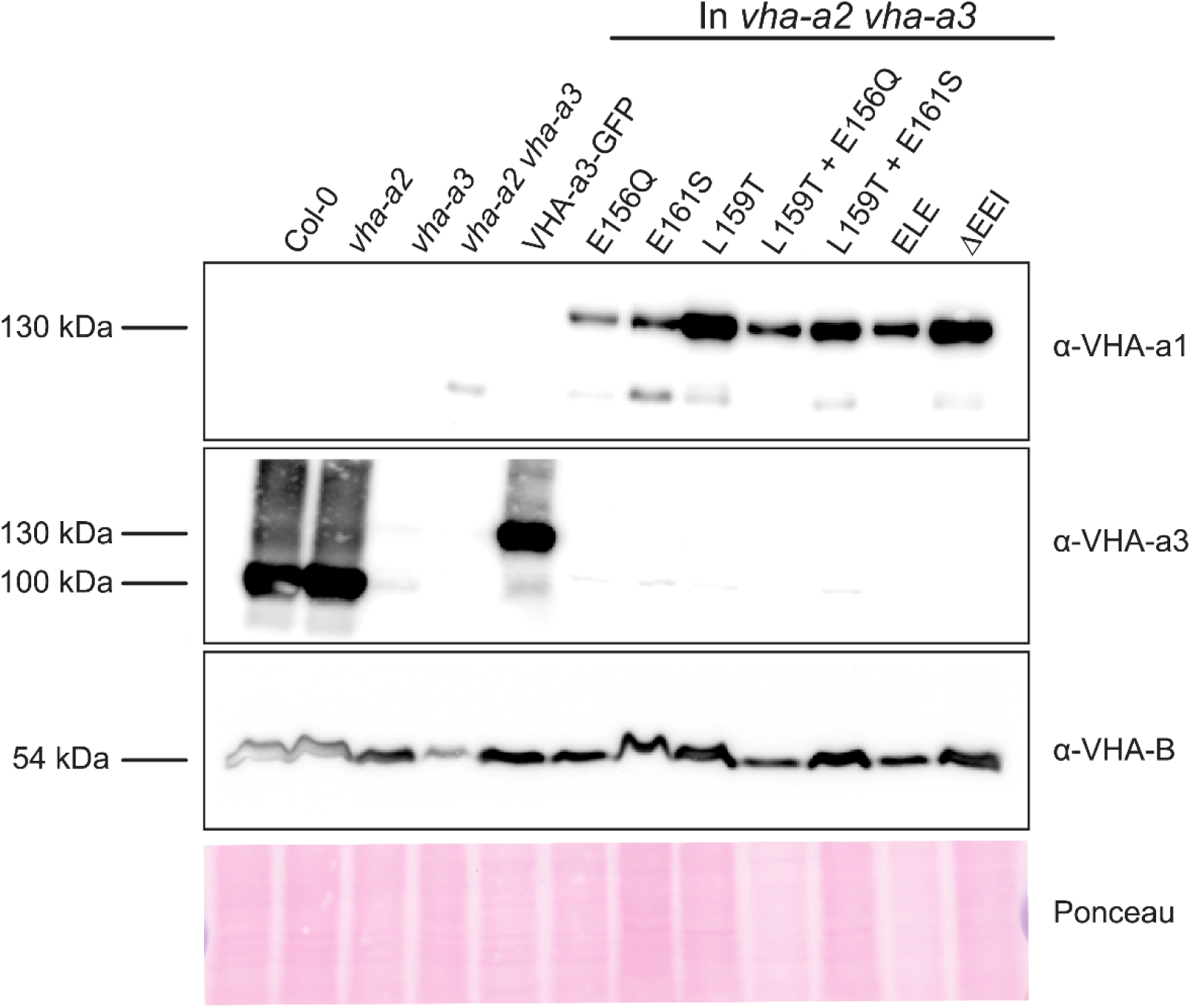
Protein levels of the mutated VHA-a proteins at the tonoplast. Abundance of the GFP tagged proteins was determined via western blot. Tonoplast membrane proteins were separated by SDS-PAGE and subsequently immunoblotted with an anti-VHA-a1, VHA-a3 and VHA-B antibodies. Protein loading is indicated by Ponceau staining of the membrane.

**Supplemental Figure 10.**
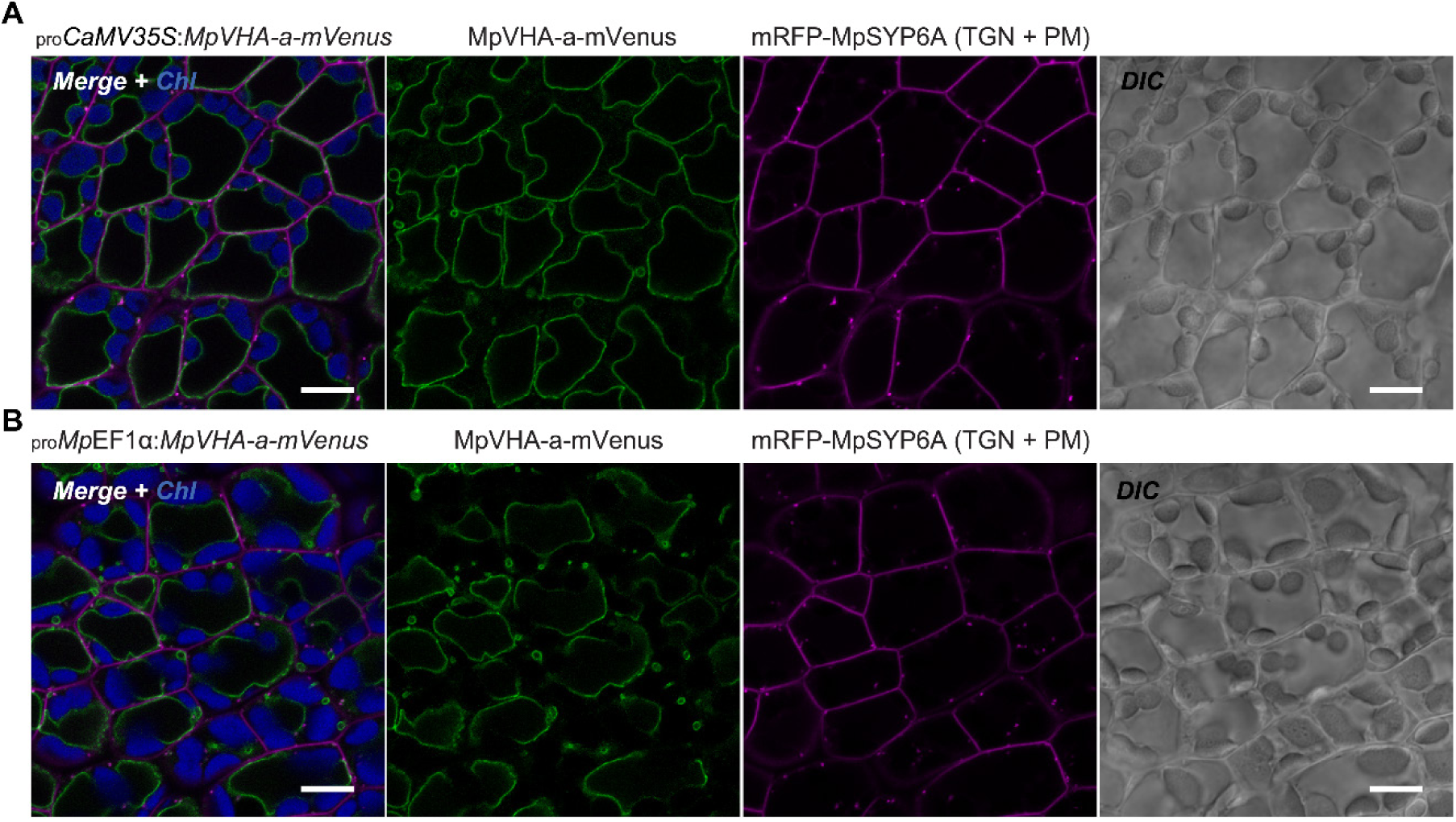
Subcellular localization of MpVHA-a-mVenus in *M*. *polymorpha*. (**A** and **B**) Single confocal images of *M*. *polymorpha* dorsal thallus cells co-expressing mRFP-MpSYP6A and MpVHA-a-mVenus driven by the CaMV35S **(A)** or Mp*EF1α* **(B)** promoter. Green, magenta, and blue pseudo colors indicate fluorescence from mVenus, mRFP, and chlorophyll, respectively. DIC images are also shown. Scale bars = 10 μm.

**Supplemental Figure 11.**
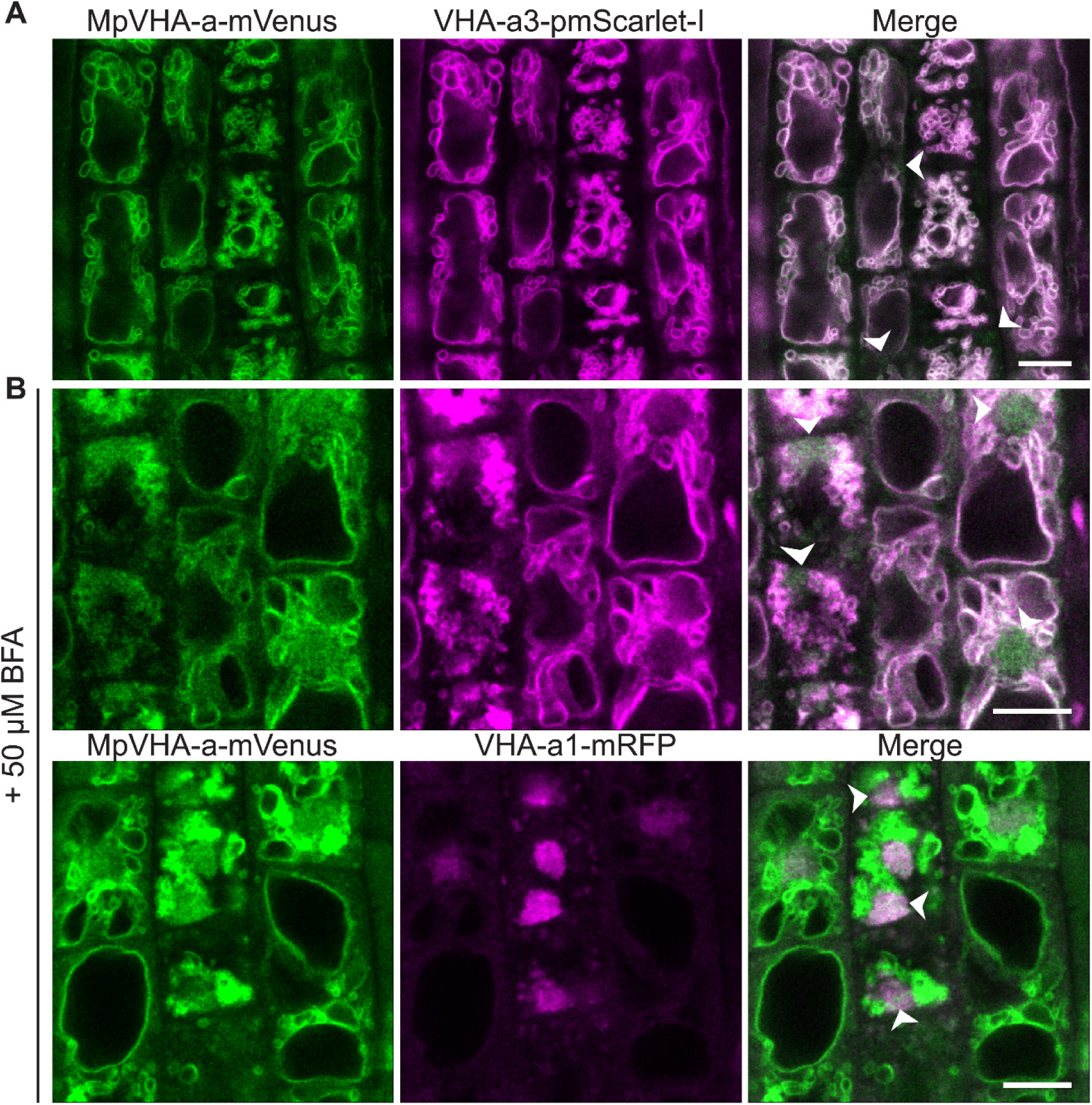
MpVHA-a-mVenus is dual localized at the TGN/EE and tonoplast in Arabidopsis wildtype root cells. *UBQ10:MpVHA-a-mvenus* was co-expressed with *UBQ10:VHA-a3-pmScarlet-I* and *VHA-a1:VHA-a1-mRFP* in Arabidopsis wildtype. Confocal analysis was done on root cells. **(A)** MpVHA-a-mvenus predominantly co-localizes with VHA-a3-pmScarlet-I at the tonoplast. **(B)** TGN localization was confirmed by treatment of root cells with 50 µm BFA for 3 hours. The core of BFA compartments were labelled with VHA-a1-mRFP and MpVHA-mvenus. Scale bars = 10 µm.

**Supplemental Figure 12.**
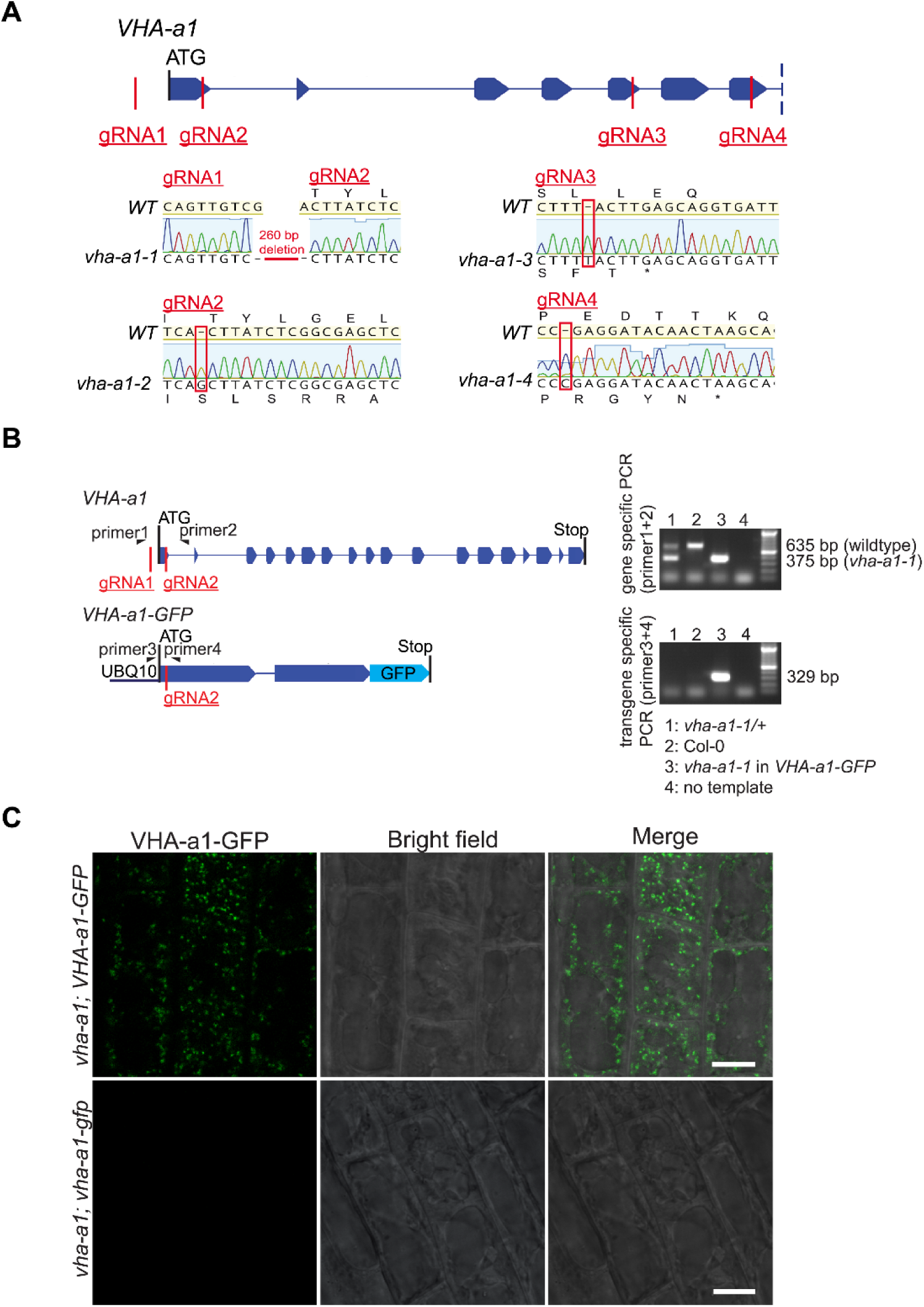
v*h*a*-a1-1* can be distinguished from the wildtype allele without sequencing and mutations in *VHA-a1-GFP* lead to absence of GFP signal. **(A)** The exon-intron structure of the first seven exons of *VHA-a1* shows the sites which were targeted in independent CRISPR approaches. *vha-a1-1*, *vha-a1-2*, *vha-a1-3* and *vha-a1-4* are examples for *vha-a1* alleles that were obtained. **(B)** Mutations in *VHA-a1* were distinguished from mutations in *VHA-a1-GFP* by use of specific primers. Shown are the exon-intron structures of gene and transgene. Primer 1 is *VHA-a1* specific, while primer 3 is *VHA-a1-GFP* specific. On an agarose gel, PCR products from *vha-a1-1* (260 bp deleted) could be distinguished from PCR products from the wildtype allele. **(C)** Root cells were analyzed by CLSM. In plants with mutations in *UBQ10:VHA-a1-GFP* corresponding to *vha-a1-2* (+1 bp at CRISPR site 2 leading to frameshift and early stop codon) no GFP signal was detected suggesting that VHA-a1-GFP was absent. Scale bars = 10 µm.

**Supplemental Figure 13.**
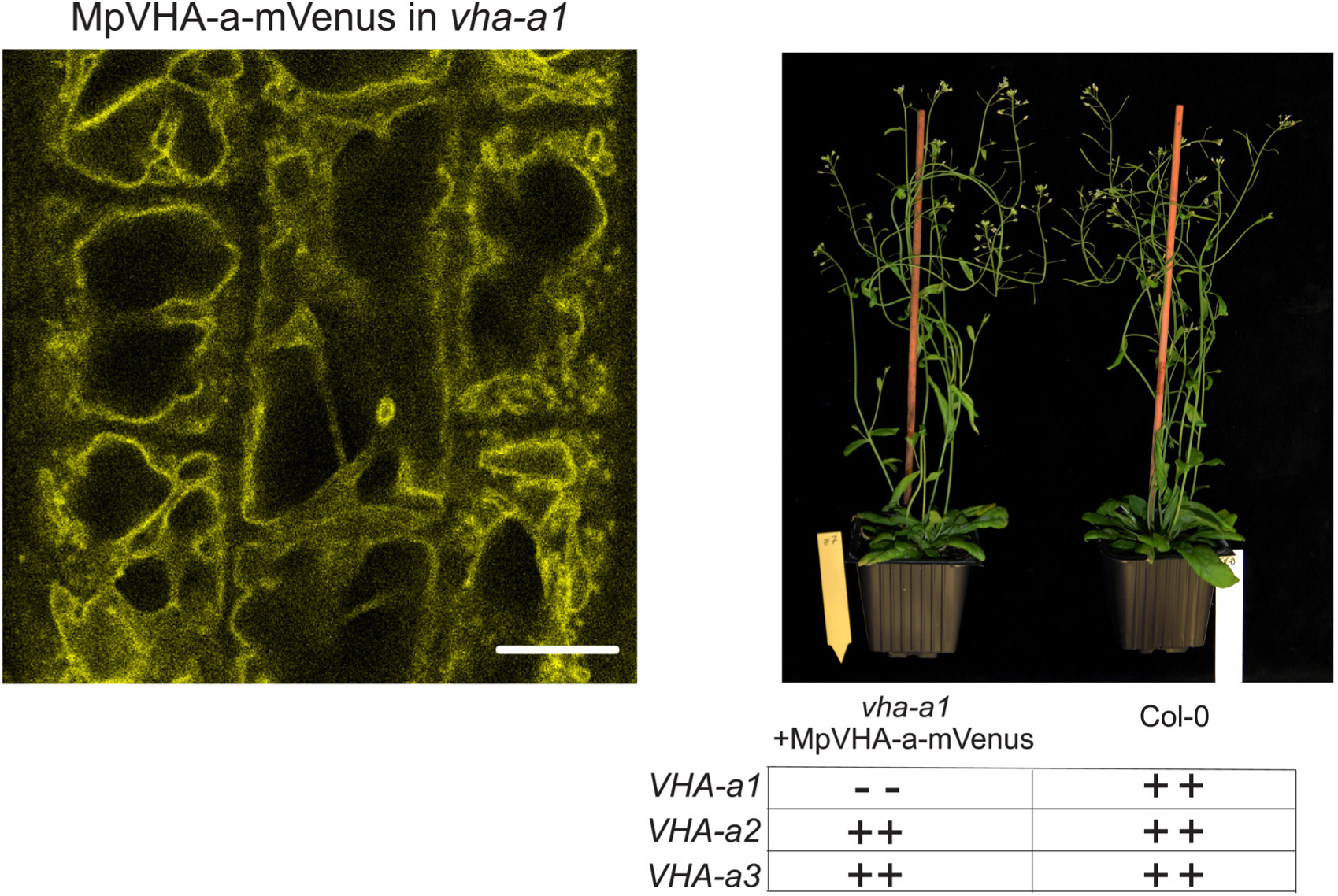
MpVHA-a-mVenus is dual localized in *vha-a1*. *vha-a1 (Cas9^+^)* was crossed with *UBQ10:MpVHA-a-mVenus* in Col-0 background and F1 seedlings were analyzed by CLSM. MpVHA-a-mVenus was dual localized at the TGN/EE and tonoplast in plants that were subsequently identified as *vha-a1* mutants. MpVHA-a-mVenus does not rescue the pollen phenotype of *vha-a1* as seen from the short siliques. Scale bar = 10 µm.

**Supplemental Figure 14.**
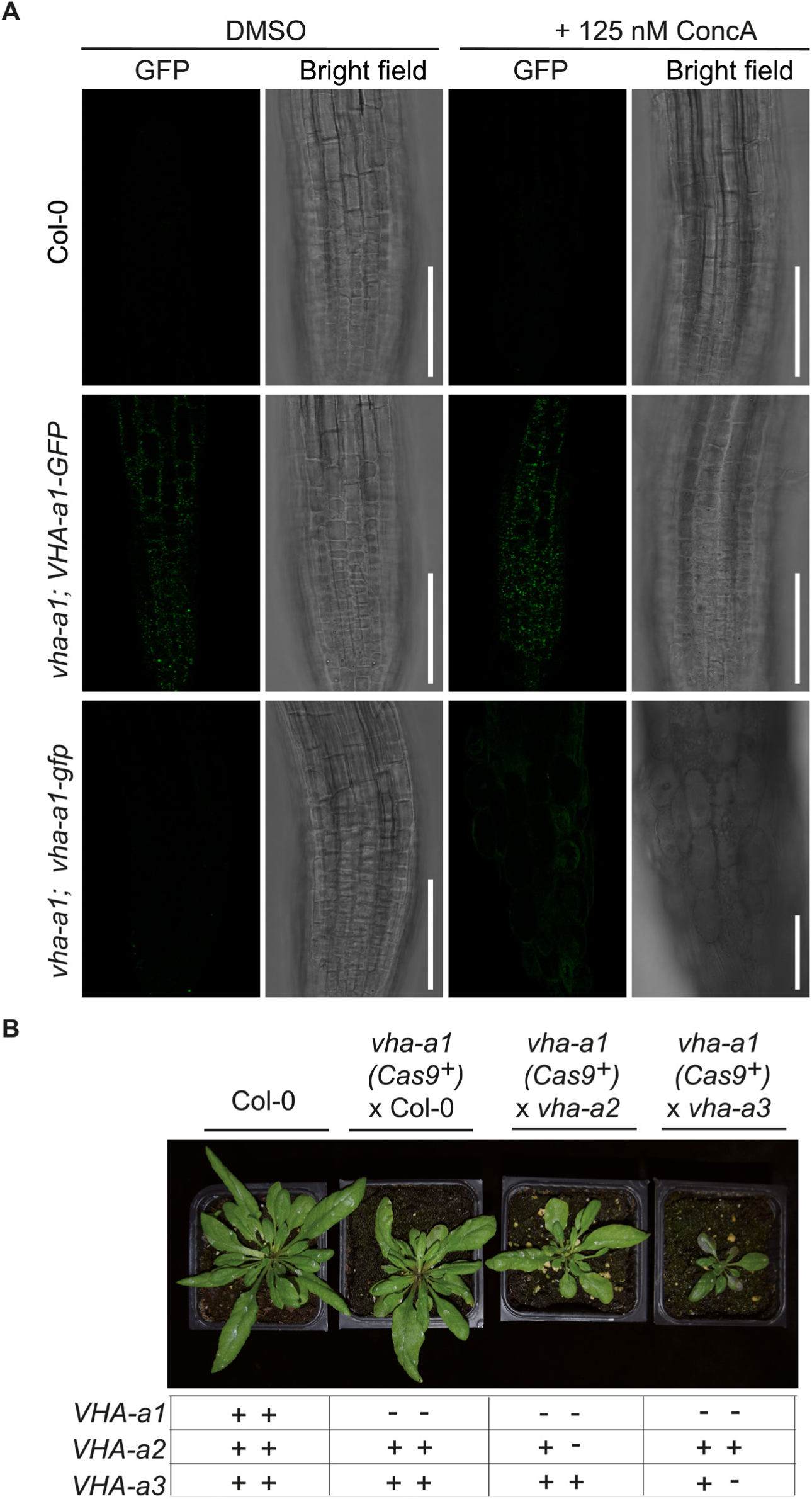
v*h*a*-a1* is hypersensitive to Concanamycin A (ConcA) and *vha-a1 vha-a2/+* and *vha-a1 vha-a3/+* have reduced rosette sizes. **(A)** Root morphology of 4-day-old etiolated seedlings. *vha-a1* roots are hypersensitive to 125 nM ConcA, in contrast to wildtype roots and roots of plants expressing *vha-a1;VHA-a1-GFP*. Scale bars = 75 µm. **(B)** *vha-a1* (*Cas9*^+^) was crossed with the *vha-a2* and the *vha-a3* single mutants. Analysis of F1 plants showed that *vha-a1 vha-a2/+* and *vha-a1 vha-a3/+* are reduced in growth. Rosettes of 5-week-old plants are shown. Plants were grown under long day conditions (22°C and 16 hours light).

## References

Alconada, A., Bauer, U., and Hoflack, B. (1996). A tyrosine-based motif and a casein kinase II phosphorylation site regulate the intracellular trafficking of the varicella- zoster virus glycoprotein I, a protein localized in the trans-Golgi network. The EMBO journal 15: 6096–6110.

Alexander, M.P. (1969). Differential Staining of Aborted and Nonaborted Pollen. Stain Technology 44: 117–122.

Barkla, B.J., Vera-Estrella, R., Maldonado-Gama, M., and Pantoja, O. (1999). Abscisic acid induction of vacuolar H+-ATPase activity in mesembryanthemum crystallinum is developmentally regulated. Plant Physiol. 120: 811–820.

Bassil, E., Ohto, M.-A., Esumi, T., Tajima, H., Zhu, Z., Cagnac, O., Belmonte, M., Peleg, Z., Yamaguchi, T., and Blumwald, E. (2011). The Arabidopsis intracellular Na+/H+ antiporters NHX5 and NHX6 are endosome associated and necessary for plant growth and development. Plant Cell 23: 224–239.

Boevink, P., Oparka, K., Santa Cruz, S., Martin, B., Betteridge, A., and Hawes, C. (1998). Stacks on tracks: the plant Golgi apparatus traffics on an actin/ER network. Plant J. 15: 441–447.

Bos, K., Wraight”, C., and Stanley, K.K. (1993). TGN38 is maintained in the trans-Golgi network by a tyrosine-containing motif in the cytoplasmic domain. The EMBO journal 12: 2219–2228.

Brüx, A., Liu, T.-Y., Krebs, M., Stierhof, Y.-D., Lohmann, J.U., Miersch, O., Wasternack, C., and Schumacher, K. (2008). Reduced V-ATPase activity in the trans-Golgi network causes oxylipin-dependent hypocotyl growth Inhibition in Arabidopsis. Plant Cell 20: 1088–1100.

Bryant, N.J. and Stevens, T.H. (1997). Two separate signals act independently to localize a yeast late Golgi membrane protein through a combination of retrieval and retention. J. Cell Biol. 136: 287–297.

Cereghino, J.L., Marcusson, E.G., and Emr, S.D. (1995). The cytoplasmic tail domain of the vacuolar protein sorting receptor Vps10p and a subset of VPS gene products regulate receptor stability, function, and localization. Mol. Biol. Cell 6: 1089–1102.

Cooper, A.A. and Stevens, T.H. (1996). Vpsl0p Cycles between the Late-Golgi and Prevacuolar Compartments in Its Function as the Sorting Receptor for Multiple Yeast Vacuolar Hydrolases. J. Cell Biol. 133.

Cotter, K., Stransky, L., McGuire, C., and Forgac, M. (2015). Recent Insights into the Structure, Regulation, and Function of the V-ATPases. Trends Biochem. Sci. 40: 611–622.

Craft, J., Samalova, M., Baroux, C., Townley, H., Martinez, A., Jepson, I., Tsiantis, M., and Moore, I. (2005). New pOp/LhG4 vectors for stringent glucocorticoid- dependent transgene expression in Arabidopsis. Plant J. 41: 899–918.

Crooks, G.E., Hon, G., Chandonia, J.-M., and Brenner, S.E. (2004). WebLogo: a sequence logo generator. Genome Res. 14: 1188–1190.

daSilva, L.L.P., Snapp, E.L., Denecke, J., Lippincott-Schwartz, J., Hawes, C., and Brandizzi, F. (2004). Endoplasmic Reticulum Export Sites and Golgi Bodies Behave as Single Mobile Secretory Units in Plant Cells. Plant Cell 16: 1753–1771.

Deschamps, A., Colinet, A.-S., Zimmermannova, O., Sychrova, H., and Morsomme, P. (2020). A new pH sensor localized in the Golgi apparatus of Saccharomyces cerevisiae reveals unexpected roles of Vph1p and Stv1p isoforms. Sci. Rep. 10: 1881.

Dettmer, J., Hong-Hermesdorf, A., Stierhof, Y.-D., and Schumacher, K. (2006). Vacuolar H+-ATPase activity is required for endocytic and secretory trafficking in Arabidopsis. Plant Cell 18: 715–730.

Dettmer, J., Schubert, D., Calvo-Weimar, O., Stierhof, Y.-D., Schmidt, R., and Schumacher, K. (2005). Essential role of the V-ATPase in male gametophyte development. Plant J. 41: 117–124.

Dragwidge, J.M., Scholl, S., Schumacher, K., and Gendall, A.R. (2019). NHX-type Na+(K+)/H+ antiporters are required for TGN/EE trafficking and endosomal ion homeostasis in Arabidopsis thaliana. J. Cell Sci. 132.

Falk, T. et al. (2019). Growing and cultivating the forest genomics database, TreeGenes. Database 2019.

Fecht-Bartenbach, J. von der, von der Fecht-Bartenbach, J., Bogner, M., Krebs, M., Stierhof, Y.-D., Schumacher, K., and Ludewig, U. (2007). Function of the anion transporter AtCLC-d in the trans-Golgi network. The Plant Journal 50: 466–474.

Finnigan, G.C., Cronan, G.E., Park, H.J., Srinivasan, S., Quiocho, F.A., and Stevens, T.H. (2012). Sorting of the Yeast Vacuolar-type, Proton-translocating ATPase Enzyme Complex (V-ATPase): IDENTIFICATION OF A NECESSARY AND SUFFICIENT GOLGI/ENDOSOMAL RETENTION SIGNAL IN Stv1p. J. Biol. Chem. 287: 19487–19500.

Finnigan, G.C., Hanson-Smith, V., Houser, B.D., Park, H.J., and Stevens, T.H. (2011). The reconstructed ancestral subunit a functions as both V-ATPase isoforms Vph1p and Stv1p in Saccharomyces cerevisiae. MBoC 22: 3176–3191.

Forgac, M. (2007). Vacuolar ATPases: rotary proton pumps in physiology and pathophysiology. Nat. Rev. Mol. Cell Biol. 8: 917–929.

Futai, M., Sun-Wada, G.-H., Wada, Y., Matsumoto, N., and Nakanishi-Matsui, M. (2019). Vacuolar-type ATPase: A proton pump to lysosomal trafficking. Proc. Jpn. Acad. Ser. B Phys. Biol. Sci. 95: 261–277.

Geldner, N., Hyman, D.L., Wang, X., Schumacher, K., and Chory, J. (2007). Endosomal signaling of plant steroid receptor kinase BRI1. Genes Dev. 21: 1598– 1602.

Goodstein, D.M., Shu, S., Howson, R., Neupane, R., Hayes, R.D., Fazo, J., Mitros, T., Dirks, W., Hellsten, U., Putnam, N., and Rokhsar, D.S. (2012). Phytozome: a comparative platform for green plant genomics. Nucleic Acids Res. 40: D1178–86.

Grefen, C., Donald, N., Hashimoto, K., Kudla, J., Schumacher, K., and Blatt, M.R. (2010). A ubiquitin-10 promoter-based vector set for fluorescent protein tagging facilitates temporal stability and native protein distribution in transient and stable expression studies. Plant J. 64: 355–365.

Hajdukiewicz, P., Svab, Z., and Maliga, P. (1994). The small, versatilepPZP family ofAgrobacterium binary vectors for plant transformation. Plant Mol. Biol. 25: 989–994.

Hanitzsch, M., Schnitzer, D., Seidel, T., Golldack, D., and Dietz, K.-J. (2007). Transcript level regulation of the vacuolar H(+)-ATPase subunit isoforms VHA-a, VHA-E and VHA-G in Arabidopsis thaliana. Mol. Membr. Biol. 24: 507–518.

Hellens, R.P., Edwards, E.A., Leyland, N.R., Bean, S., and Mullineaux, P.M. (2000). pGreen: a versatile and flexible binary Ti vector for Agrobacterium-mediated plant transformation. Plant Mol. Biol. 42: 819–832.

Hillmer, S., Viotti, C., and Robinson, D.G. (2012). An improved procedure for low-temperature embedding of high-pressure frozen and freeze-substituted plant tissues resulting in excellent structural preservation and contrast. J. Microsc. 247: 43–47.

Ishizaki, K., Chiyoda, S., Yamato, K.T., and Kohchi, T. (2008). Agrobacterium-mediated transformation of the haploid liverwort Marchantia polymorpha L., an emerging model for plant biology. Plant Cell Physiol. 49: 1084–1091.

Ishizaki, K., Nishihama, R., Ueda, M., Inoue, K., Ishida, S., Nishimura, Y., Shikanai, T., and Kohchi, T. (2015). Development of Gateway Binary Vector Series with Four Different Selection Markers for the Liverwort Marchantia polymorpha. PLoS One 10: e0138876.

Kanazawa, T. et al. (2016). SNARE Molecules in Marchantia polymorpha: Unique and Conserved Features of the Membrane Fusion Machinery. Plant Cell Physiol. 57: 307– 324.

Kawasaki-Nishi, S., Bowers, K., Nishi, T., Forgac, M., and Stevens, T.H. (2001a). The amino-terminal domain of the vacuolar proton-translocating ATPase a subunit controls targeting and in vivo dissociation, and the carboxyl-terminal domain affects coupling of proton transport and ATP hydrolysis. J. Biol. Chem. 276: 47411–47420.

Kawasaki-Nishi, S., Nishi, T., and Forgac, M. (2001b). Yeast V-ATPase complexes containing different isoforms of the 100-kDa a-subunit differ in coupling efficiency and in vivo dissociation. J. Biol. Chem. 276: 17941–17948.

Kozlov, A.M., Darriba, D., Flouri, T., Morel, B., and Stamatakis, A. (2019). RAxML-NG: a fast, scalable and user-friendly tool for maximum likelihood phylogenetic inference. Bioinformatics 35: 4453–4455.

Krebs, M., Beyhl, D., Görlich, E., Al-Rasheid, K.A.S., Marten, I., Stierhof, Y.-D., Hedrich, R., and Schumacher, K. (2010). Arabidopsis V-ATPase activity at the tonoplast is required for efficient nutrient storage but not for sodium accumulation. Proc. Natl. Acad. Sci. U. S. A. 107: 3251–3256.

Krebs, M., Held, K., Binder, A., Hashimoto, K., Den Herder, G., Parniske, M., Kudla, J., and Schumacher, K. (2012). FRET-based genetically encoded sensors allow high-resolution live cell imaging of Ca2 dynamics. The Plant Journal 69: 181–192.

Kriegel, A., Andrés, Z., Medzihradszky, A., Krüger, F., Scholl, S., Delang, S., Patir- Nebioglu, M.G., Gute, G., Yang, H., Murphy, A.S., Peer, W.A., Pfeiffer, A., Krebs, M., Lohmann, J.U., and Schumacher, K. (2015). Job Sharing in the Endomembrane System: Vacuolar Acidification Requires the Combined Activity of V- ATPase and V-PPase. Plant Cell 27: 3383–3396.

Kubota, A., Ishizaki, K., Hosaka, M., and Kohchi, T. (2013). Efficient Agrobacterium- mediated transformation of the liverwort Marchantia polymorpha using regenerating thalli. Biosci. Biotechnol. Biochem. 77: 167–172.

Labun, K., Montague, T.G., Gagnon, J.A., Thyme, S.B., and Valen, E. (2016). CHOPCHOP v2: a web tool for the next generation of CRISPR genome engineering. Nucleic Acids Res. 44: W272–6.

Lampropoulos, A., Sutikovic, Z., Wenzl, C., Maegele, I., Lohmann, J.U., and Forner, J. (2013). GreenGate---a novel, versatile, and efficient cloning system for plant transgenesis. PLoS One 8: e83043.

Lanfear, R., Frandsen, P.B., Wright, A.M., Senfeld, T., and Calcott, B. (2017). PartitionFinder 2: New Methods for Selecting Partitioned Models of Evolution for Molecular and Morphological Phylogenetic Analyses. Mol. Biol. Evol. 34: 772–773.

Leidi, E.O., Barragán, V., Rubio, L., El-Hamdaoui, A., Ruiz, M.T., Cubero, B., Fernández, J.A., Bressan, R.A., Hasegawa, P.M., Quintero, F.J., and Pardo, J.M. (2010). The AtNHX1 exchanger mediates potassium compartmentation in vacuoles of transgenic tomato. Plant J. 61: 495–506.

Leng, X.H., Manolson, M.F., and Forgac, M. (1998). Function of the COOH-terminal domain of Vph1p in activity and assembly of the yeast V-ATPase. J. Biol. Chem. 273: 6717–6723.

Luo, Y. et al. (2015). V-ATPase activity in the TGN/EE is required for exocytosis and recycling in Arabidopsis. Nat Plants 1: 15094.

Madeira, F., Park, Y.M., Lee, J., Buso, N., Gur, T., Madhusoodanan, N., Basutkar, P., Tivey, A.R.N., Potter, S.C., Finn, R.D., and Lopez, R. (2019). The EMBL-EBI search and sequence analysis tools APIs in 2019. Nucleic Acids Res. 47: W636– W641.

Marshansky, V., Rubinstein, J.L., and Grüber, G. (2014). Eukaryotic V-ATPase: novel structural findings and functional insights. Biochim. Biophys. Acta 1837: 857–879.

Moore, I., Gälweiler, L., Grosskopf, D., Schell, J., and Palme, K. (1998). A transcription activation system for regulated gene expression in transgenic plants. Proc. Natl. Acad. Sci. U. S. A. 95: 376–381.

Nebenführ, A., Ritzenthaler, C., and Robinson, D.G. (2002). Brefeldin A: deciphering an enigmatic inhibitor of secretion. Plant Physiol. 130: 1102–1108.

Neubert, C., Graham, L.A., Black-Maier, E.W., Coonrod, E.M., Liu, T.-Y., Stierhof, Y.- D., Seidel, T., Stevens, T.H., and Schumacher, K. (2008). Arabidopsis has two functional orthologs of the yeast V-ATPase assembly factor Vma21p. Traffic 9: 1618– 1628.

Nothwehr, S.F., Roberts, C.J., and Stevens, T.H. (1993). Membrane protein retention in the yeast Golgi apparatus: dipeptidyl aminopeptidase A is retained by a cytoplasmic signal containing aromatic residues. J. Cell Biol. 121: 1197–1209.

Rienmüller, F., Dreyer, I., Schönknecht, G., Schulz, A., Schumacher, K., Nagy, R., Martinoia, E., Marten, I., and Hedrich, R. (2012). Luminal and cytosolic pH feedback on proton pump activity and ATP affinity of V-type ATPase from Arabidopsis. J. Biol. Chem. 287: 8986–8993.

Roth, R. et al. (2018). A rice Serine/Threonine receptor-like kinase regulates arbuscular mycorrhizal symbiosis at the peri-arbuscular membrane. Nat. Commun. 9: 4677.

Roy, A., Kucukural, A., and Zhang, Y. (2010). I-TASSER: a unified platform for automated protein structure and function prediction. Nat. Protoc. 5: 725–738.

Samalova, M., Brzobohaty, B., and Moore, I. (2005). pOp6/LhGR: a stringently regulated and highly responsive dexamethasone-inducible gene expression system for tobacco. Plant J. 41: 919–935.

Schäfer, W., Stroh, A., Berghöfer, S., Seiler, J., Vey, M., Kruse, M.L., Kern, H.F., Klenk, H.D., and Garten, W. (1995). Two independent targeting signals in the cytoplasmic domain determine trans-Golgi network localization and endosomal trafficking of the proprotein convertase furin. The EMBO journal 14: 2424–2435.

Scheuring, D., Viotti, C., Krüger, F., Künzl, F., Sturm, S., Bubeck, J., Hillmer, S., Frigerio, L., Robinson, D.G., Pimpl, P., Schumacher, K., and Ku, F. (2011). Multivesicular Bodies Mature from the Trans-Golgi Network/Early Endosome in Arabidopsis. Plant Cell 23: 1–20.

Schürholz, A.-K.K., López-Salmerón, V., Li, Z., Forner, J., Wenzl, C., Gaillochet, C., Augustin, S., Vilches Barro, A., Fuchs, M., Gebert, M., Lohmann, J.U., Greb, T. and Wolf, S. (2018). A comprehensive toolkit for inducible, cell type-specific gene expression in Arabidopsis. Plant Physiol. 178: 40–53.

Smyth, D.R., Bowman, J.L., and Meyerowitz, E.M. (1990). Early flower development in Arabidopsis. Plant Cell 2: 755–767.

Stemmer, M., Thumberger, T., Del Sol Keyer, M., Wittbrodt, J., and Mateo, J.L. (2015). CCTop: An Intuitive, Flexible and Reliable CRISPR/Cas9 Target Prediction Tool. PLoS One 10: e0124633.

Sze, H., Li, X., and Palmgren, M.G. (1999). Energization of plant cell membranes by H+- pumping ATPases. Regulation and biosynthesis. Plant Cell 11: 677–690.

Sze, H., Schumacher, K., Müller, M.L., Padmanaban, S., and Taiz, L. (2002). A simple nomenclature for a complex proton pump: VHA genes encode the vacuolar H(+)- ATPase. Trends Plant Sci. 7: 157–161.

Vasanthakumar, T., Bueler, S.A., Wu, D., Beilsten-Edmands, V., Robinson, C.V., and Rubinstein, J.L. (2019). Structural comparison of the vacuolar and Golgi V-ATPases from Saccharomyces cerevisiae. Proc. Natl. Acad. Sci. U. S. A. 116: 7272–7277.

Viotti, C., Bubeck, J., Stierhof, Y.-D., Krebs, M., Langhans, M., van den Berg, W., van Dongen, W., Richter, S., Geldner, N., Takano, J., Jürgens, G., de Vries, S.C., Robinson, D.G., and Schumacher, K. (2010). Endocytic and secretory traffic in Arabidopsis merge in the trans-Golgi network/early endosome, an independent and highly dynamic organelle. Plant Cell 22: 1344–1357.

Viotti, C., Krüger, F., Krebs, M., Neubert, C., Fink, F., Lupanga, U., Scheuring, D., Boutté, Y., Frescatada-Rosa, M., Wolfenstetter, S., Sauer, N., Hillmer, S., Grebe, M., and Schumacher, K. (2013). The endoplasmic reticulum is the main membrane source for biogenesis of the lytic vacuole in Arabidopsis. Plant Cell 25: 3434–3449.

Waadt, R., Krebs, M., Kudla, J., and Schumacher, K. (2017). Multiparameter imaging of calcium and abscisic acid and high-resolution quantitative calcium measurements using R-GECO1-mTurquoise in Arabidopsis. New Phytol. 216: 303–320.

Wang, Z.-P., Xing, H.-L., Dong, L., Zhang, H.-Y., Han, C.-Y., Wang, X.-C., and Chen, Q.-J. (2015). Egg cell-specific promoter-controlled CRISPR/Cas9 efficiently generates homozygous mutants for multiple target genes in Arabidopsis in a single generation. Genome Biol. 16: 144.

Ward, J.M., Reinders, A., Hsu, H.-T., and Sze, H. (1992). Dissociation and Reassembly of the Vacuolar H-ATPase Complex from Oat Roots. PLANT PHYSIOLOGY 99: 161–169.

Wassmer, T., Kissmehl, R., Cohen, J., and Plattner, H. (2006). Seventeen a-Subunit Isoforms of Paramecium V-ATPase Provide High Specialization in Localization and Function. Molecular Biology of the Cell 17: 917–930.

Wegrzyn, J.L., Lee, J.M., Tearse, B.R., and Neale, D.B. (2008). TreeGenes: A forest tree genome database. Int. J. Plant Genomics 2008: 412875.

Wilcox, C.A., Redding, K., Wright, R., and Fuller, R.S. (1992). Mutation of a tyrosine localization signal in the cytosolic tail of yeast Kex2 protease disrupts Golgi retention and results in default transport to the vacuole. Mol. Biol. Cell 3: 1353–1371.

Xiang, Y., Molloy, S.S., Thomas, L., and Thomas, G. (2000). The PC6B cytoplasmic domain contains two acidic clusters that direct sorting to distinct trans-Golgi network/endosomal compartments. Mol. Biol. Cell 11: 1257–1273.

Zhao, J., Benlekbir, S., and Rubinstein, J.L. (2015). Electron cryomicroscopy observation of rotational states in a eukaryotic V-ATPase. Nature 521: 241–245.

